# *Dscam* gene triplication causes neocortical overinhibition in Down syndrome

**DOI:** 10.1101/2020.01.03.894113

**Authors:** H Liu, RN Caballero-Florán, T Yang, JM Hull, G Pan, R Li, MW Veling, LL Isom, KY Kwan, ZJ Huang, PG Fuerst, PM Jenkins, B Ye

## Abstract

A growing number of molecules have been identified as regulators of inhibitory synapse development, but whether dysregulated expression of these molecules contribute to brain disorders is poorly understood. Here we show that Down syndrome cell adhesion molecule (*Dscam*) regulates the inhibition of neocortical pyramidal neurons (PyNs) in a level-dependent fashion. Loss of *Dscam* impairs inhibitory neuron development and function. In the Ts65Dn mouse model for Down syndrome, where *Dscam* is overexpressed, GABAergic innervation of cortical PyNs by chandelier and basket cells is increased. Genetic normalization of *Dscam* expression rescues the excessive GABAergic innervation and the increased inhibition of PyNs. These findings demonstrate excessive GABAergic innervation and inhibition in the neocortex of Down syndrome mouse model and identify *Dscam* overexpression as the cause. They also implicate dysregulated *Dscam* levels as a potential pathogenic driver in related neurological disorders.

## Introduction

GABAergic neurons mediate synaptic inhibition in the mammalian brain (1, 2), and altered GABAergic inhibition has emerged as a nexus of circuit dysfunction in a number of neurodevelopmental disorders (3, 4). A growing number of molecules have been identified in GABAergic synapse formation and synaptic transmission (4–9). Intriguingly, copy number variations of these genes are commonly found in different neurological diseases (7), suggesting that dysregulated expression of these genes might be a pathogenic driver. However, whether or not this is true remains largely unknown.

Down syndrome cell adhesion molecule (*Dscam*) is an evolutionarily conserved type I transmembrane protein (10). In human, *Dscam* gene is localized in the Down syndrome critical region on chromosome 21 (hCh21) (10). We previously showed in *Drosophila* that *Dscam* protein levels determine the sizes of presynaptic terminals without requiring the ectodomain diversity of the *Drosophila Dscam* gene (11). Moreover, others reported that overexpression of *Dscam* impairs synaptic targeting and transmission in *Drosophila* (12, 13). These findings suggest that dysregulated *Dscam* levels might contribute to neuronal defects in brain disorders. In fact, altered *Dscam* expression levels have been reported in multiple brain disorders, including Down syndrome (DS) (14), autism spectrum disorders (ASD) (15–17), intractable epilepsy (18), bipolar disorder (19), and possibly Fragile X syndrome (11, 12, 20, 21). Although recent findings suggest a conserved role of *Dscam* in promoting presynaptic growth in vertebrates (22, 23), whether dysregulated *Dscam* expression results in neuronal defects in brain disorders remains unknown.

In this work, we sought to determine the effects of altered expression levels of *Dscam* in mouse models of DS, in which the *Dscam* gene is triplicated. Previous studies have shown that enhanced GABAergic inhibition impairs cognition in Ts65Dn mice (24–28), the most widely used DS animal model (29, 30). A series of studies have demonstrated excessive GABAergic inhibition in the hippocampus of Ts65Dn mice (27, 31–39). However, whether GABAergic signaling is altered in other brain regions and, if it is, what genes on hCh21 cause it, are poorly understood. Here, by combining recently developed genetic tools for sparse labeling and whole-cell patch-clamp, we found excessive GABAergic innervation and inhibition of PyNs by chandelier (ChC) and basket cells in the neocortex of Ts65Dn mice. Genetic normalization of *Dscam* expression levels rescued the presynaptic overgrowth and excessive synaptic transmission of GABAergic neurons. Consistently, loss of *Dscam* impaired ChC presynaptic terminals and reduced GABAergic inhibition of neocortical PyNs. In addition, we found that ChC axon terminal growth and synaptogenesis are coupled and positively correlated with neocortical *Dscam* levels in both wide-type and Ts65Dn mice. These findings uncover novel molecular, cellular and synaptic mechanisms for DS brain disorders and highlight the critical role of *Dscam* levels in regulating neocortical inhibition. Dysregulated *Dscam* expression levels may be a common contributor to GABAergic dysfunctions in associated neurological diseases.

## Materials and Methods

### Mice breeding and tamoxifen administration

Age-matched littermates were used for all experiments. Ts65Dn (Ts(17(16))65Dn) was purchased from Jackson Laboratory (Stock No: 005252) and maintained by crossing with C57BL/C3H F1 hybrid males. For labeling single ChCs in Ts65Dn mice, Ts65Dn females were crossed with Nkx2.1-CreER+/−, Ai14−/−, *Dscam*+/− male mice of mixed C57BL/C3H background to obtain euploid, Ts65Dn and Ts65Dn: *Dscam*+/+/− littermates with Nkx2.1-CreER+/−, Ai14+/− transgenes. Tamoxifen (Sigma, T5648-1G), dissolved in corn oil, was delivered to P0 pups by intraperitoneal injection at the dosage of 80 mg/kg body weight. Pseudodam was prepared in advance to lactate the pups as the pups were often discarded by Ts65Dn dam after the drug delivery. For labeling single ChCs in *Dscam*2j mice, female *Dscam*+/− mice of C3H background were crossed with Nkx2.1-CreER+/−, Ai14−/−, *Dscam*+/− male mice of mixed C57BL/C3H background to obtain *Dscam*+/− and *Dscam*- pups with Nkx2.1-CreER+/−, Ai14+/− transgenes. Tamoxifen was delivered to P0 pups as above. Strong pups were sacrificed at P0 to keep the litter size 4-5 to increase the surviving rate of *Dscam*- pups. Nkx2.1-CreER+/−, Ai14−/−, *Dscam*+/− male mice were removed from the cage before female laboring as the male adults is likely to attack and sacrifice *Dscam*-pups.

For electrophysiology recordings, Ts65Dn females were crossed with C3H *Dscam*+/− male mice to obtain euploid, Ts65Dn and Ts65Dn: *Dscam*+/+/− littermates. C3H *Dscam*+/− mice were interbred to obtain *Dscam*+/− and *Dscam*- pups for recordings.

PCR genotyping was performed on purified tail tips at least 2 times to confirm the genotype according to the protocol by Jackson Laboratory. All work involving mice was approved by the University of Michigan Institutional Animal Care and Use Committee (IACUC) and in accordance with the National Research Council Guide for the Care and Use of Laboratory Animals (NIH).

### Tissue processing

P28 mice were euthanized by CO2 and immediately followed by intracardial infusion of 4% paraformaldehyde (PFA) with 4% sucrose in 1x PBS. The brains were then removed and post-fixed in 4 % PFA overnight at 4 °C, followed by incubation in 30% sucrose (wt/vol) for at least 1 day. The brains were embedded in the OCT compound (Fisher HealthCare), frozen in −20 °C overnight and then sectioned to 100 µm-thick slices with a Leica CM3050S cryostat. Sectioned brain slices were kept in 1x PBS containing 0.05% sodium azide at 4 °C until immunostaining.

### Immunohistochemistry

For ChC and AIS staining, coronal brain sections (100 μm) were blocked with 8% BSA in PBST (1x PBS + 0.1% Triton x-100) containing 0.05 % sodium azide at 37°C for 1 hour, and then incubated with the following primary antibodies in the blocking solution at 37°C overnight: anti-mCherry (AB0081; 1:300; SICGEN) for neuronal morphology and anti-phospho-Iϰβ-α (14D4 rabbit monoclonal antibody; 1:500; Cell Signaling) for labeling the AIS. After washing 3 time (1 hour each time) at 37°C in PBST, brain slices were incubated with the following secondary antibodies in the blocking solution at 37°C overnight: donkey anti-goat-rhodamine RX (RRX) (705-297-003; 1:300; Jackson ImmunoResearch), donkey anti-rabbit-AlexaFluor 488 (711-545-152; 1:300; Jackson ImmunoResearch). After washing 3x for a total of 1 hour at 37°C in PBST, the slices were mounted in sRIMS (40).

The procedure for staining of GABAergic neurons and PyNs was the same as above, except that antibody incubations were done at room temperature and different primary antibodies were used. Primary antibodies used were anti-parvalbumin (PVG213; 1:1,000, Swant Inc), anti-CAMKIIα (6G9; 1:5,000, LSBio). Secondary antibodies used were donkey anti-goat-RRX (705-297-003; 1:500; Jackson ImmunoResearch) and donkey anti-mouse-AlexaFluor 488 (715-545-150; 1:500; Jackson ImmunoResearch).

For staining perisomatic GABAergic synapses, antigen retrieval was conducted in 10 μM sodium citrate for 20 min at 95°C (41). After a brief rinse in PBS, the slices were blocked in blocking buffer (1x PBS, 0.3% Triton x-100, 3% normal donkey serum, 0.05 % sodium azide) for 1 hour at room temperature (RT), and then incubated with the following primary antibodies in the blocking solution at 4°C overnight: mouse anti-Bassoon (SAP7F407; 1:1,000; ENZO), G.P. anti-VGAT (131 004; 1:1,000; Synaptic System), rabbit anti-GRASP1 (1:2,000) (42). After washing 3 time (20 mins each time) at RT in PBST, brain slices were incubated with the following secondary antibodies in the blocking solution at RT 3 hours: donkey anti-mouse-AlexaFluor 488 (715-545-150; 1:300; Jackson ImmunoResearch), donkey anti-G.P.-RRX (706-295-148; 1:300; Jackson ImmunoResearch), donkey anti-rabbit-AlexaFluor 647 (711-605-152; 1:300; Jackson ImmunoResearch),. The slices were mounted for imaging after washing 3x for a total of 1 hour at RT in PBST.

The procedure for c-fos staining was as described previously (43). Briefly, brain sections were washed three times (10-min each) with 1x PBS, and then treated sequentially with the following solutions: 2% H2O2, 0.3% NaOH/1x PBS for 20 min, 1x PBS for 5 min twice, 0.3% glycine/1x PBS for 10 min, 1x PBS for 5 min twice, 0.03% SDS/1x PBS for 10 min, and 1x PBS for 5 min twice. The brain sections were then blocked in 3% normal donkey serum/0.1% Triton X-100/1x PBS for 1 hr, and subsequently incubated overnight in blocking solution that contained anti-c-Fos antibody (#2250; 1:2000 cell signaling). The sections were washed in 1x PBS for three 20-min washes, and incubated with biotinylated secondary antibody (711-066-152; 1:200; Jackson Immunoresearch) in blocking solution for 2 hours. After washing in 1x PBS for three 10-min washes, the sections were processed for ABC amplification (PK-6100; 1:500 in PBS dilution; Vector Laboratories) for 1 hour, washed in 1x PBS for 10 min three times, incubated in 0.05% diaminobenzidine (DAB)/0.015% H2O2/1x PBS for 3 min, and then washed in 1x PBS for 10 min three times.

### Image acquisition and quantitative analysis

All images were acquired from layer II/III of the ACC in the mouse neocortex by using a Leica SP5 confocal microscope equipped with a resonant scanner, except the samples for perisomatic GABAergic synapses (see below). For ChCs imaging, different fluorescence channels were imaged sequentially with the pinhole set at airy 1. A 63x objective lens with a numerical aperture of 1.4 was used. Confocal image stacks were collected with 100 continuous optical sections at 0.3-μm z-steps. The cell body was positioned around the middle of the 30-μm in depth. Before quantification, the image stacks were maximally projected along the z-axis. A region of 120 μm (length) × 80 μm (width) with the cell body in the top middle was set for quantification. Cartridges and boutons that colocalized with AIS were quantified by the NeuroLucida software (MBF Bioscience). Cartridge/bouton number was defined as the number of cartridges/boutons within this region. Cartridge length was defined as the distance from the first bouton that colocalized with AIS to the last one colocalized with AIS in that cartridge. Bouton size was defined as the length of the bouton in parallel to the AIS. We quantified the sizes of boutons in the 10 cartridges nearest to the cell body. Interbouton distance was defined as the distance between two neighboring boutons.

Samples for perisomatic GABAergic synapses were imaged on a Leica SP8 confocal microscope with a 63x objective lens (NA 1.4). For each PyN, a single confocal image was taken at the z-position where PyN cell body occupied the most area. Images were deconvoluted with the Huygens software (Scientific Volume Imaging). Perisomatic VGAT+ and Bassoon+ puncta were quantified i n d efined Py N so ma regions, as shown in Figure 6B, by the NeuroLucida software (MBF Bioscience).

For c-fos and parvabumin imaging, a 20x objective lens with a NA of 1.4 was used. The z-steps were 1 μm. Five optical sections were maximally projected before quantification. Three fields were imaged for each mouse. Cells were counted in a defined area (252.4 μm × 200 μm) for each field manually with the assistance of the NeuroLucida software. To eliminate experimenter’s bias, these experiments were carried out in double-blind fashion. Two double-blind methods were used. First, the images acquired by the primary experimenter (H.L.) was coded and randomized by the second lab member (R.L. or M.V.). After the primary experimenter quantified the data, the data were decoded for statistical analysis. ChCs from 2 littermates were quantified in this method. Second, the mouse brains from the primary experimenter (H.L.) were coded and randomized by the second lab member (R.L. or M.V.). The second lab member sent the encoded mouse brains to the third lab member (T.Y.) for sectioning. After the primary experimenter immunostained and quantified the encoded brain sections, the data were decoded for a final statistical analysis. ChCs from 3 littermates and the c-fos, parvabumin and basket cell synapse quantifications were performed in this method.

### Western blotting

Mouse neocortices were removed immediately after PFA perfusion. Similar volumes of tissues were homogenized in SDS sample buffer and loaded for electrophoresis. The proteins were transferred to PVDF membranes and blocked with 5% milk (wt/vol) for 1 h at room temperature. The blots were incubated overnight at 4 °C with goat anti *Dscam* (AF3666, 1:500, R&D systems) and mouse anti-tubulin (12G10, 1:5,000, DSHB). After washing, blots were incubated for 2 h with HRP-conjugated secondary antibodies and developed by chemiluminescence (Catalog# 32106, Pierce). Quantification was performed using ImageJ.

### Electrophysiology recordings and analysis

Electrophysiological recordings of spontaneous and miniature inhibitory postsynaptic currents were performed as described previously (44). Brains were obtained from euploid, Ts65Dn, and Ts65Dn/*Dscam*+/+/− mice at around P28. The animals were decapitated under isoflurane and USP anesthesia, and the brain was quickly removed from the skull and placed in 4°C slicing solution containing (in mM) 62.5 NaCl, 2.5 KCl, 1.25 KH2PO4, 26 NaHCO3, 5 MgCl2, 0.5CaCl2, 20 glucose and 100 sucrose (pH maintained at 7.4 by saturation with 95% O2 + 5%CO2). Coronal brain slices (300-350 μm thick) containing layers II/III of the ACC neocortex were cut with a microtome (VF-300, Compresstome). The slices were then transferred to a holding chamber and maintained at room temperature in artificial cerebrospinal fluid (ACSF) containing (in mM) 125 NaCl, 2.9 KCl, 1.25 KH2PO4, 26 NaHCO3, 1 MgCl2, 2 CaCl2 and 20 glucose, pH 7.4 (with 95% O2 and 5% CO2 bubbling through the solution) for at least 1 hour prior to recording. After equilibration, individual slices were transferred to the recording chamber continuously perfused with ACSF (1-2 mL/min). Recording micropipettes were pulled from borosilicate glass capillaries (1.5 mm O.D. Harvard Apparatus) for a final resistance of 3-6 MΩ and filled with a solution containing (in mM) 135 CsCl, 4 NaCl, mM GTP, Mg-ATP, 0.5 CaCl2, 5 EGTA and 10 HEPES. The signals were recorded with an Axoclamp 700B amplifier (Axon Instruments, Union City, CA). Voltage clamp recordings were obtained from neurons in layers II/III of the ACC region; the cells in these brain regions were identified using a Nikon Eclipse FN-1 microscope with a 40x water-immersion objective and a DAGE MTI IR-1000 video camera. Neurons were visualized using IR-DIC to evaluate their orientation and morphology. Whole-cell patch-clamp recordings for cells with a high cell resistance (greater than 8 GΩ before break-in). sIPSC and mIPSC recordings in voltage-clamp configuration were acquired at 2 kHz fixing the voltage at −70 mV. The IPSCs were recorded in the presence of the NMDA receptor antagonist DL-2-amino-5-phosphonopentanoic acid (AP-5) at 50 mM and the AMPA/kainite antagonist 6-cyano-7-nitroquinoxaline-2,3-dione (CNQX) at 10 μM. For measurement of mIPSCs, 1 μM tetrodotoxin (TTX) was added to the perfusion solution to block synaptic responses dependent on the AP. Access resistance was monitored throughout the experiment and experiments were canceled if changes greater than 20% occurred. Peak events were identified automatically using Minianalysis (Synaptosoft Inc.) and visually monitored to exclude erroneous noise. The frequency, amplitude and distribution of events were analyzed. One euploid neuron (1 out of 20) with mIPSC frequency 8.9 Hz was defined as an outlier by the Grubbs’ test and removed from quantification. Mean values were compared using the Student’s t-test. Data are presented as mean ± SEM.

### Statistical analysis

Data are presented as mean ± SEM. Comparisons of mean differences between groups were performed by one way ANOVA followed by Student’s t-test. P < 0.05 was considered to be statistically significant. For all quantification, *: p < 0.05; **: p < 0.01; ***: p < 0.001; ns: not significant (p > 0.05).

## Results

### Loss of *Dscam* impairs ChC presynaptic development in mice

To determine whether *Dscam* regulates GABAergic neuron development in the neocortex, we focused the analysis on ChCs for two reasons. First, ChCs are thought to be the most potent inhibitory neurons in the neocortex (45). Each ChC innervates roughly two hundred PyNs at their axon initial segments (AIS) (46), where action potentials are generated (47, 48). Second, the morphology of ChCs is relative stereotypical and reliably quantifiable (49–52). We specifically label single ChCs in the neocortex b y crossing Nkx2.1-CreER mouse line with the tdTomato reporter line Ai14 (53)(Figure 1A). A single ChC extends only a few dendritic branches but several hundreds of presynaptic terminals, called axonal cartridges, each of which innervates one or two PyNs (54). The axonal cartridges and presynaptic boutons of single ChCs were quantified i n a threedimensional volume defined based on the position of ChC cell body (Figure S1). AIS-colocalized cartridges and presynaptic boutons were quantified, as off-target varicosities lack presynaptic markers and symmetric synaptic densities (55). The AIS was immunolabeled by an anti-phospho-IϰB antibody (pIϰB) (50, 51, 53, 56), which strongly correlates with the AIS labeled by ankyrin-G antibodies in the mouse neocortex (57)(Figure S2). Quantification w as p erformed in a double-blinded fashion to avoid experimenters’ bias. The ChCs in the cortical layer II and III of the anterior cingulate cortex (ACC) were analyzed as the morphology of ChCs in this region is most stereotypical. Mice were analyzed on postnatal day 28 (P28), a time after ChC development is complete (54). Similar to its role in presynaptic development in *Drosophila*, loss of *Dscam* significantly impeded the development of ChC presynaptic terminals (Figures 1B-E). Compared to heterozygous littermates, the number and length of individual cartridges were significantly reduced in *Dscam*-- mice by 8% and 12%, respectively so that the total cartridge length for each ChC was reduced by 23% (Figures 1C-E).

**Fig. 1.**
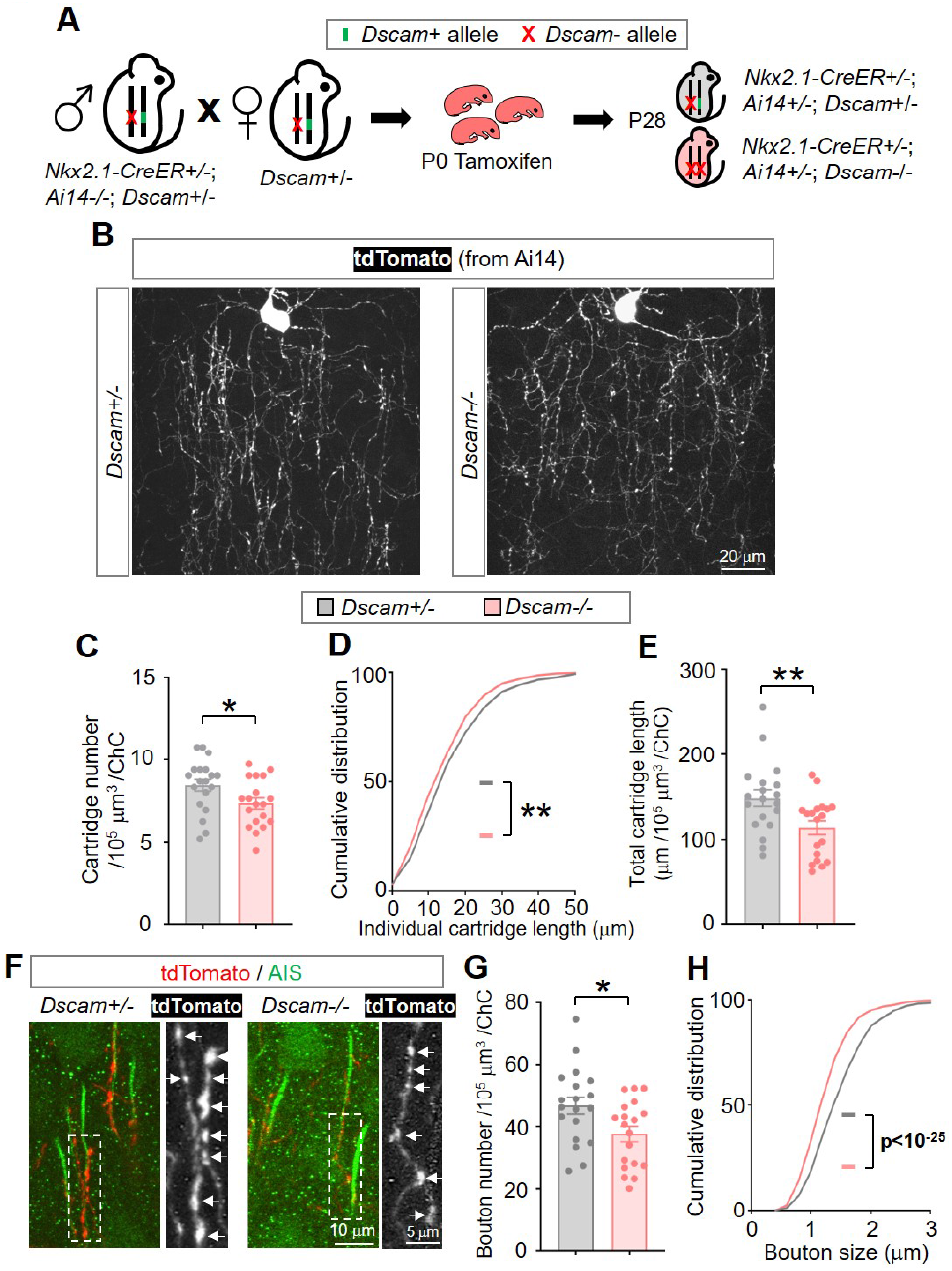
Loss of *Dscam* impairs the growth of ChC axon cartridges and boutons. (A) A schematic of the procedure that produced the mice for the experiments. (B) Representative images of single ChCs in layer II/III of anterior cingulate cortex (ACC). Shown are ChCs of *Dscam*2j/+ (+/−, left) and *Dscam*2j/2j (−/−, right) mice at P28. All ChC images in this paper are from this brain region of P28 mice. Scale bar, 20 μm. (C-E) Quantification of axon cartridge number (C), length (D) and the total cartridge length (E). For each ChC, all axon cartridges (15-40) innervating the AIS of PyNs in a volume of 120 μm (length) × 80 μm (width) × 30 μm (thickness) with the ChC cell body in the top middle were analyzed. 4-6 ChCs were analyzed for each mouse, and 4 *Dscam*+/− and 4 *Dscam*- mice were analyzed. Each dot in (C) and (E) represents one ChC. Sample numbers in (B) and (D) are 19 for *Dscam*+/− and 19 for *Dscam*-. In (D), cumulative plots show the distributions of cartridge length. Sample numbers are 462 for *Dscam*+/− and 402 for *Dscam*-.. Unless specified, mean ± SEM is shown in the figures, and the statistical tests are one-way ANOVA followed by Student’s t test. *: p < 0.05; **: p < 0.01; ***: p < 0.001; ns: not significant (p > 0.05). (F) Representative images of ChC axon cartridges innervating the AIS of PyNs. Cartridges of single ChCs were labeled with tdTomato (Red). The AIS of PyNs were labeled by anti-phospho-IκB (pIκB, Green). The arrows point to presynaptic boutons of ChCs in the boxed regions. (G-H) Quantification of bouton number (G) and size (H). For each ChC, all boutons (38 194) in axon cartridges that innervate AIS in the defined volume (Figure S1) were analyzed for bouton numbers; boutons (31 84) in the 10 cartridges nearest to the cell body were analyzed for bouton sizes. 4-6 ChCs were analyzed in each mouse. 4 *Dscam*+/− and 4 *Dscam*- mice were analyzed. In (G), each dot represents one ChC, N: 19 for *Dscam*+/− and *Dscam*-. In (H), accumulative plots show the distributions of bouton size. N: 1079 for *Dscam*+/− and 1004 for *Dscam*-.

Moreover, the numbers and sizes of presynaptic boutons were significantly reduced by 20% and 16%, respectively (Figures 1F-H). Interestingly, no difference was observed in the interbouton distance between these two groups (Figure S3A), indicating that *Dscam* does not regulate bouton density. These results demonstrate that *Dscam* is required for ChC presynaptic development in mice.

Loss of *Dscam* impaired GABAergic inhibition of PyNs To determine whether defective GABAergic synaptogenesis caused by loss of *Dscam* impairs GABAergic synaptic transmission, whole-cell patch-clamp was applied to record PyNs in layers II/III of the ACC from acute neocortical brain slices. Consistent with impaired axonal growth and synaptogenesis in ChCs, we found that the average frequency of miniature inhibitory postsynaptic currents (mIPSCs) was 35% less in *Dscam*- mice than that in heterozygous littermates (Figures 2A-B). In addition, the average amplitude of mIPSCs was 42% less, ¬suggesting that postsynaptic responses were impaired by loss of *Dscam* (Figures 2A and C). Similar changes were observed in the frequency and amplitude of spontaneous IPSC (sIPSCs) (Figures 2D-F). Consistent with the reduced GABAergic transmission, we observed hyperactivity in neocortical PyNs, as indicated by c-fos expression levels (58). There was 5-fold more c-fos-positive PyNs in *Dscam*- than in *Dscam*+/− mice (Figures 2G-H). Taken together, these data suggest that *Dscam* is required for the normal level of inhibition of PyNs in the neocortex.

**Fig. 2.**
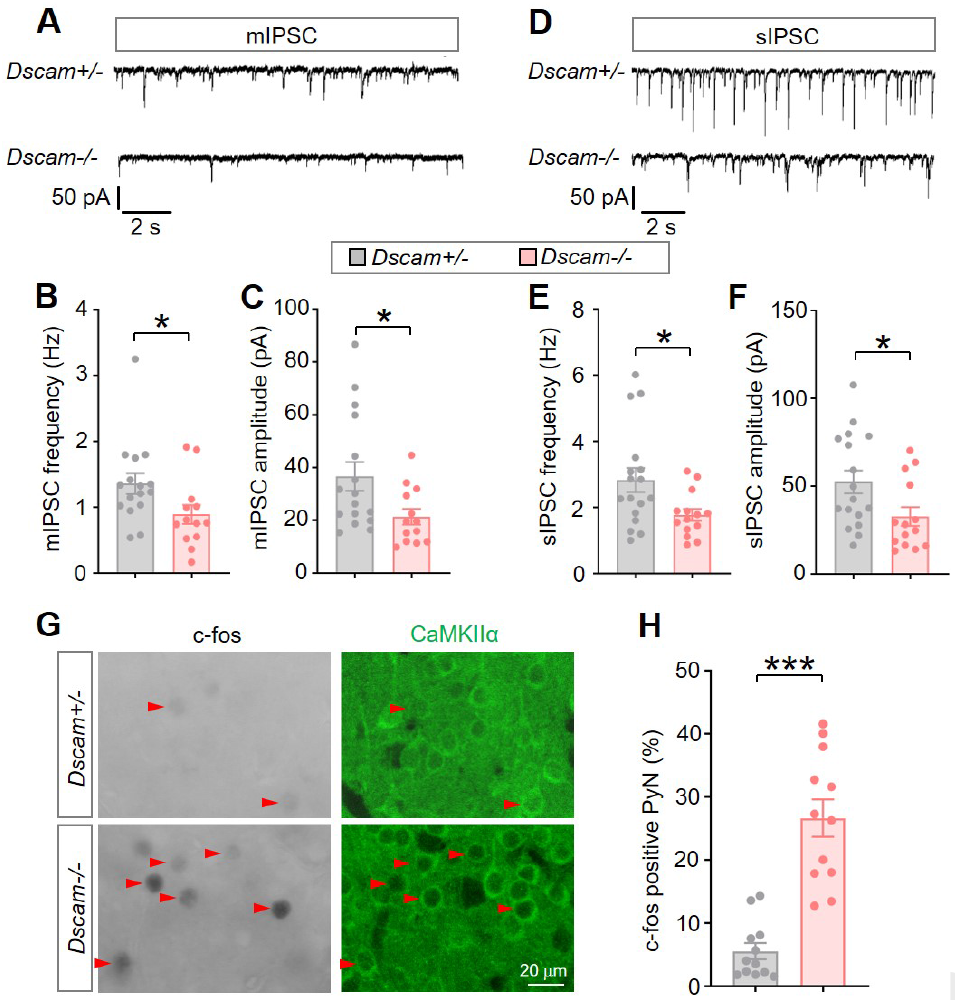
Loss of *Dscam* impairs GABAergic inhibition of neocortical PyNs. (A) Representative traces of mIPSCs from PyNs in layer II/III of ACC in *Dscam*+/− and *Dscam−/−* brain slices. (B-C) Quantification of mIPSC frequency (B) and amplitude (C). 3-5 PyNs were recorded for each mouse. 5 *Dscam*+/− and 4 *Dscam−/−* mice were analyzed. N: 16 for *Dscam*+/−, and 13 for *Dscam−/−*. (D) Representative traces of sIPSCs from PyNs in layer II/III of ACC in *Dscam*+/− and *Dscam−/−* brain slices. (E-F) Quantification of sIPSC frequency (E) and amplitude (F). 3 5 PyNs were recorded for each mouse. 5 *Dscam*+/− and 4 *Dscam−/−* mice were analyzed. N: 17 for *Dscam*+/−, 14 for *Dscam−/−*. (G) Representative images of c-fos levels in *Dscam*+/− and *Dscam−/−* mice. C-fos levels reflect neuronal activity. Anti-CaMKIIα immunostaining marks PyNs. The red arrowheads point to the PyNs with detectable c-fos immunostaining. Images were acquired from layer II/III of the ACC at P28. (H) c-fos levels are increased in *Dscam−/−* mice. The graph shows the percentage of the percentage of c-fos-positive PyNs in total PyNs. Each dot represents the value in one randomly selected field for imaging. Three fields were imaged in each mouse. 4 *Dscam*+/− and 4 *Dscam−/−* mice were analyzed.

ChCs exhibit excessive presynaptic terminals and boutons in Ts65Dn mice Next, we determined whether the presynaptic development of ChCs is altered in the Ts65Dn mouse model of DS, where *Dscam* is present in 3 copies (29) and overexpressed (Figure S4). Compared to euploid littermates, the number and length of individual axonal cartridges were significantly increased in Ts65Dn mice by 28% and 11%, respectively (Figures 3A-C).

**Fig. 3.**
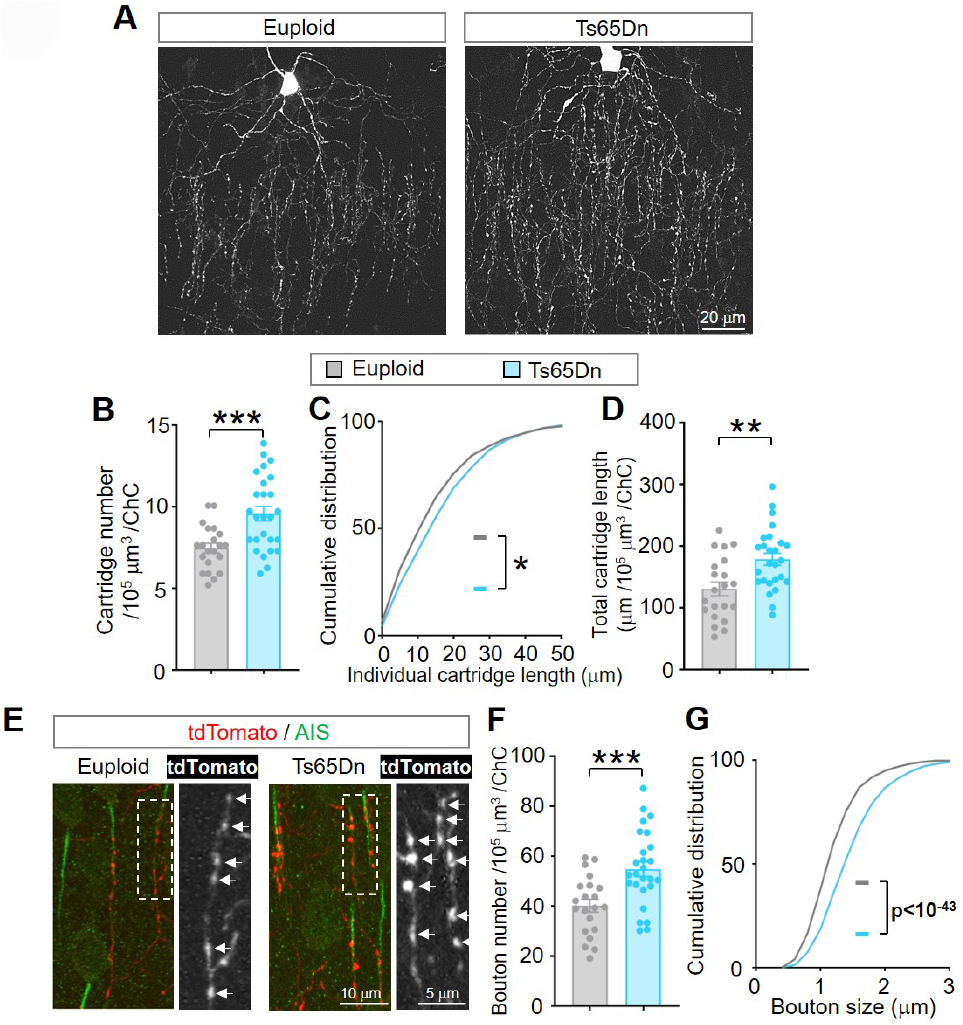
ChC axonal terminals and presynaptic boutons are overgrown in Ts65Dn mice. (A) Representative images of single ChCs in wild-type (euploid, left) and Ts65Dn (right) mice at P28. Scale bar, 20 μm. (B-D) Quantification of axon cartridge number (B), length (C) and the total cartridge length (D). Each dot in (B) and (D) represents one ChC. Sample number (n): 21 for euploid and 26 for Ts65Dn. In (C), cumulative plots show the distributions of cartridge length, n=452 for euploid, 717 for Ts65Dn. Sample numbers are shown in the figure. (E) Representative images of ChC axon cartridges innervating the AIS of PyNs. Cartridges of single ChCs were labeled with tdTomato. The AIS of PyNs were labeled by anti-phospho-IκB (pIκB). The arrows point to presynaptic boutons of ChCs in the boxed regions. (F-G) Quantification of bouton number (F) and size (G). 4-6 ChCs were analyzed for each mouse. 4 euploid and 5 Ts65Dn mice were analyzed. In (F), each dot represents one ChC, N: 21 for euploid and 26 for Ts65Dn. In (G), accumulative plots show the distributions of bouton size, n=1149 for euploid, 1655 for Ts65Dn.

The total cartridge length, which is the sum of individual cartridges in the quantified volume, was increased by 37% (Figure 3D). In addition, ChC showed a significant increase in both the number and size of synaptic boutons in Ts65Dn mice. The bouton number for each ChC is increased by 36% (Figures 3E-F). The average size of presynaptic boutons was enlarged by 21% (Figures 3E and G). The increased ChC presynaptic bouton number and size are consistent with previous reports of increased number of the puncta positive of vesicular GABA transporter proteins (VGAT) and enlarged inhibitory synapses in the neocortex of Ts65Dn mice (59, 60). The average interbouton distance between neighboring boutons showed a significant, though subtle, decrease in the trisomy mice (Figure S3B), suggesting a slight increase in bouton density. Altogether, by labeling and quantifying single interneurons in the ACC, we found excessive presynaptic terminals and boutons in Ts65Dn mice, which is opposite of what we observed in *Dscam*- mice.

Normalization of *Dscam* levels rescues ChC presynaptic overgrowth in Ts65Dn mice *Dscam* is overexpressed in both the brains of DS patients and those of Ts65Dn mouse model of DS (14)(Figure S4). Previous studies in *Drosophila* have demonstrated that increased levels of *Dscam* lead to excessive growth of presynaptic terminals (11, 61). To determine whether the overexpressed *Dscam* in Ts65Dn mice causes presynaptic overgrowth in ChCs, we normalized *Dscam* gene dosage in Ts65Dn mice by crossing female Ts65Dn mice with male *Dscam*+/− mice to obtain the Ts65Dn:*Dscam*+/+/− genotype (Figure 4A). This genetic scheme also yielded regular Ts65Dn littermates (Ts65Dn:*Dscam*+/+/+). In Ts65Dn mice with 2 copies of functional *Dscam* genes, the average levels of *Dscam* proteins were statistically indistinguishable from the euploid mice (Figure 4B). Normalizing *Dscam* levels rescued ChC presynaptic overgrowth in Ts65Dn mice. Compared with Ts65Dn littermates, the ChC cartridge number, individual cartridge length and total cartridge length were reversed to levels indistinguishable from euploid in Ts65Dn:*Dscam*+/+/− mice (Figures 4C-F). In addition, the increased bouton number and bouton size were mostly rescued by normalizing *Dscam* expression (Figures 4G-I), while no change was observed in the interbouton distance (Figure S3B). These results demonstrate that ChC presynaptic terminal overgrowth is mainly caused by *Dscam* overexpression in Ts65Dn mice.

**Fig. 4.**
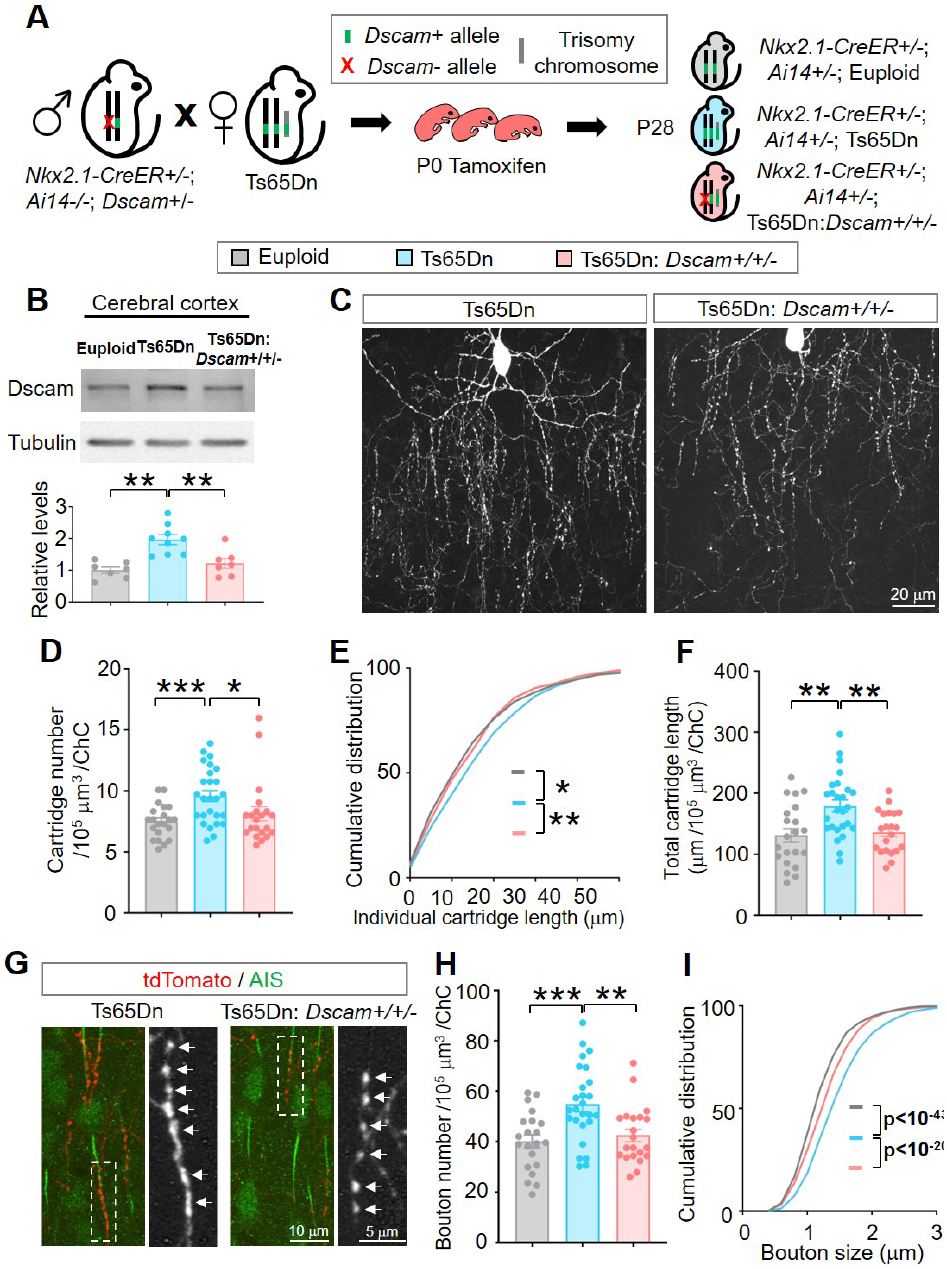
Normalizing *Dscam* expression rescues the overgrowth of ChC axon cartridges and presynaptic boutons in Ts65Dn mice. (A) A schematic of the procedure that produced the mice for the experiments. (B) *Dscam* overexpression is normalized to the euploid level in Ts65Dn mice by introducing one *Dscam* mutant allele. Shown are representative western blots (top) and quantifications (bottom) of neocortical samples from euploid, Ts65Dn, and Ts65Dn:*Dscam*+/+/−. Sample numbers are indicated in the bars. (C) Representative images of single ChCs in layer II/III of the ACC. Shown are ChCs of Ts65Dn (left) and Ts65Dn with *Dscam* allele normalized (Ts65Dn:*Dscam*+/+/−, right) mice at P28. Scale bar, 20 μm. (D-F) Quantification of axon cartridge number (D), length (E) and the total cartridge length (F). 4-6 ChCs were analyzed for each mouse; 4 euploid, 5 Ts65Dn, and 4 Ts65Dn:*Dscam*+/+/− mice were analyzed. Sample numbers are 21 for euploid, 26 for Ts65Dn, and 21 for Ts65Dn:*Dscam*+/+/− in (D) and (F). In (E), n=452 for euploid, 717 for Ts65Dn and 491 for Ts65Dn:*Dscam*+/+/−. (G) Representative images of ChC cartridges innervating the AIS of PyNs. The arrows point to presynaptic boutons of ChCs in the boxed region. (H-I) Quantification of bouton number (H) and size (I). 4-6 ChCs were analyzed for each mouse, and 4 euploid, 5 Ts65Dn, and 4 Ts65Dn:*Dscam*+/+/− mice were analyzed. In (H), each dot represents one ChC, N: 21 for euploid, 26 for Ts65Dn, and 21 for Ts65Dn:*Dscam*+/+/−. In (I), accumulative plots show the distributions of bouton size, n=1149 for euploid, 1655 for Ts65Dn and 1209 for Ts65Dn:*Dscam*+/+/−.

ChC axon terminal growth and synaptogenesis are positively coupled Presynaptic terminal growth and synaptogenesis are concurrent processes for forming proper neuronal connections (62). Little is known about whether and how these two events are orchestrated in mammalian GABAergic interneurons. The sparse labeling of ChCs offers an opportunity to address these questions at the single-cell resolution. We examined the relationship between cartridge length and three morphological aspects of synaptic boutons, namely bouton number, size, and density (as reflected by the interbouton distance), in single ChCs. We found a strong correlation between the cartridge length and the bouton number of each ChC in both wild-type (euploid) (R2 = 0.82, p < 10-7) and Ts65Dn mice (R2 = 0.79, p < 10-15) (Figure 5A). Although *Dscam* positively regulated both cartridge length and bouton number (Fig. 1e, g and Fig. 4f, h), loss of *Dscam* did not impair their coupling (R2 = 0.89, p < 10-8) (Figure 5B). There was also a significant correlation between cartridge length and bouton size in both wild-type and Ts65Dn mice (Figure 5C), though it was weaker than that between cartridge length and bouton number. Loss of *Dscam* seemed to mildly impair the coupling between cartridge length and bouton size (Figure 5D). There was no correlation between cartridge length and bouton density in any of the genotypes tested (Figures 5E and F). Taken together, these results suggest that presynaptic terminal growth and synaptogenesis are positively coupled in ChC development.

**Fig. 5.**
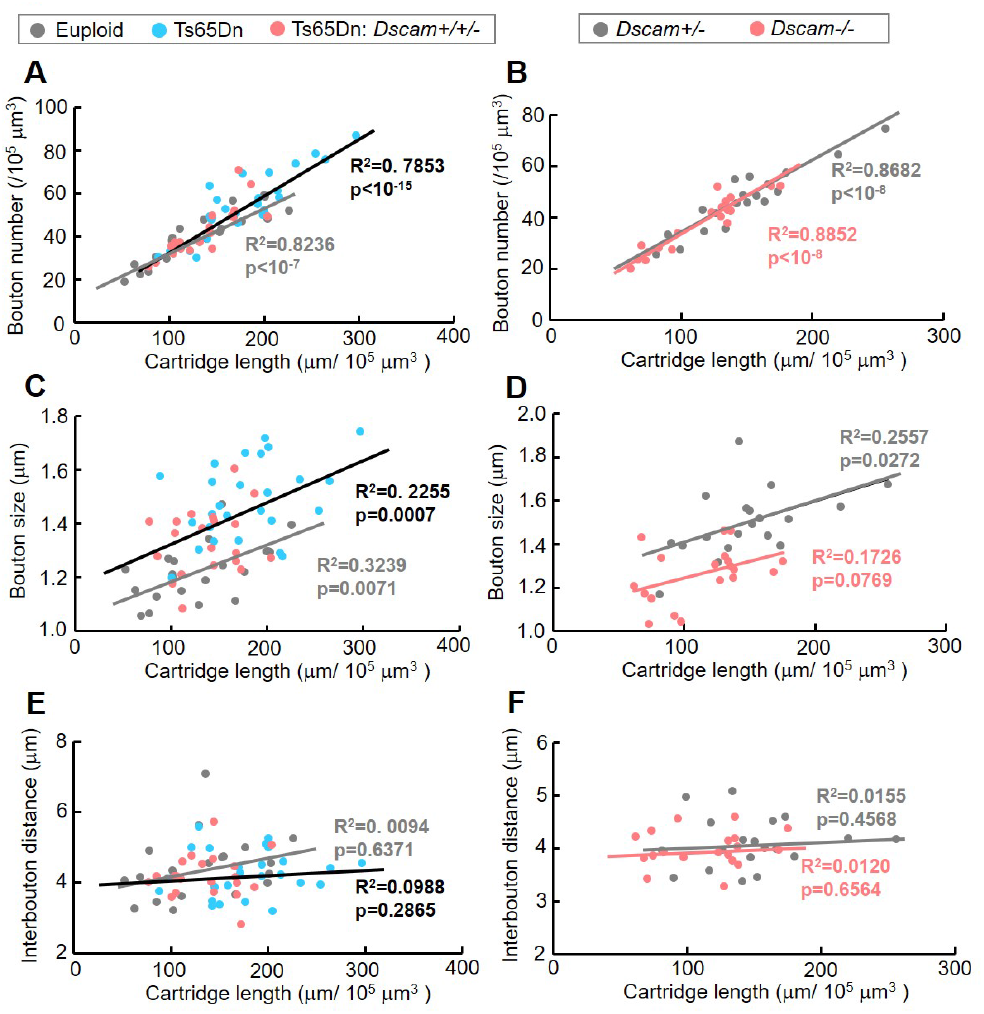
Coupling between presynaptic terminal growth and synaptogenesis in ChC development. (A) ChC cartridge length and bouton number are strongly correlated in both euploid and Ts65Dn genetic backgrounds. Each dot represents one ChC. 4-6 ChCs were analyzed for each mouse. 4 euploid, 5 Ts65Dn, 4 Ts65Dn:*Dscam*+/+/− mice were analyzed. n=21 for euploid (gray dots), 26 for Ts65Dn (blue dots), 21 Ts65Dn:*Dscam*+/+/− (coral dots). R2 and p are calculated for linear regression. The gray line indicates the trend line for gray dots, while the black line is that for blue and coral dots. (B) ChC cartridge length and bouton number are strongly correlated in *Dscam*+/− and *Dscam−/−* mice. Each dot presents one ChC. 4–6 ChCs were analyzed for each mouse. 4 *Dscam*+/− and 4 *Dscam−/−* mice were analyzed. N: 19 for *Dscam*+/− (gray dots) and 19 for *Dscam−/−* (coral dots). (C) ChC cartridge length and bouton size show weak, yet significant, correlation in euploid and Ts65Dn background. Each dot presents one ChC. (D) The correlation between cartridge length and bouton number is impaired in *Dscam−/−* mice. Each dot presents one ChC. R2 is small in both *Dscam*+/− and *Dscam−/−* mice, suggesting that linear regression only explains a small fraction of the samples. The correction is insignificant in *Dscam−/−* mice (p > 0.05). (E-F) ChC cartridge length and interbouton distance shows no significant c orrelation b etween a ny t wo g enotypes tested. Each dot presents one ChC.

ChC presynaptic cartridge length and synaptogenesis are proportional to neocortical *Dscam* levels In *Drosophila*, *Dscam* levels determine the size of presynaptic arbors (11). Western blotting results suggested that *Dscam* levels in the neocortex exhibited individual variations in euploid, Ts65Dn, and Ts65Dn:*Dscam*+/+/− mice (Figure 4B and S S5A). We plotted the neocortical *Dscam* level of each mouse, as assayed by western blotting, against the average cartridge length, bouton number, size or density in each mouse. We observed a strong correlation between *Dscam* levels and cartridge lengths, bouton number and bouton sizes in mice (Figures S5B-D). In contrast, no correlation was found between *Dscam* levels and interbouton distance (Figure S5E), again supporting that bouton density is not regulated by *Dscam* (Figures S3A-B). The dosage-dependence highlights the importance of *Dscam* expression levels in regulating ChC presynaptic development.

*Dscam* overexpression increases the number of GABAergic synapses on PyN somas in Ts65Dn mice *Dscam* is expressed in different subtypes of GABAergic neurons (63). Does its overexpression in Ts65Dn mice cause developmental defects in other types of GABAergic neurons? Basket cells are the most common cortical interneurons that preferentially innervate the somas of PyNs (1, 2). Because basket cell morphology is highly heterogeneous (64), we analyzed the GABAergic synapses formed by basket cells on PyN soma (i.e., perisomatic synapses), rather than quantifying the morphology of single basket cells as we did on ChCs. To visualize perisomatic GABAergic synapses, brain sections were triply labeled with anti-Bassoon (for presynaptic active zone) (65), anti-VGAT (for presynaptic GABAergic boutons) (66), and anti-GRASP1 (for PyN soma and proximal dendrites) (42). Most Bassoon+ puncta around a PyN soma were surrounded by or overlapped with VGAT signals (Figure S6A). We found that the average number of GABAergic synapses around each PyN soma—as indicated by Bassoon+ puncta that were either apposed to or overlapped with VGAT+ signals—was 20% more in Ts65dn neocortices than euploid controls (Figures 6A-B). Strikingly, normalizing *Dscam* expression level completely rescued the enhanced synaptogenesis (Figures 6B-D). These data suggest that the overexpressed *Dscam* in Ts65Dn mice also increases the number of GABAergic synapses on PyN somas.

**Fig. 6.**
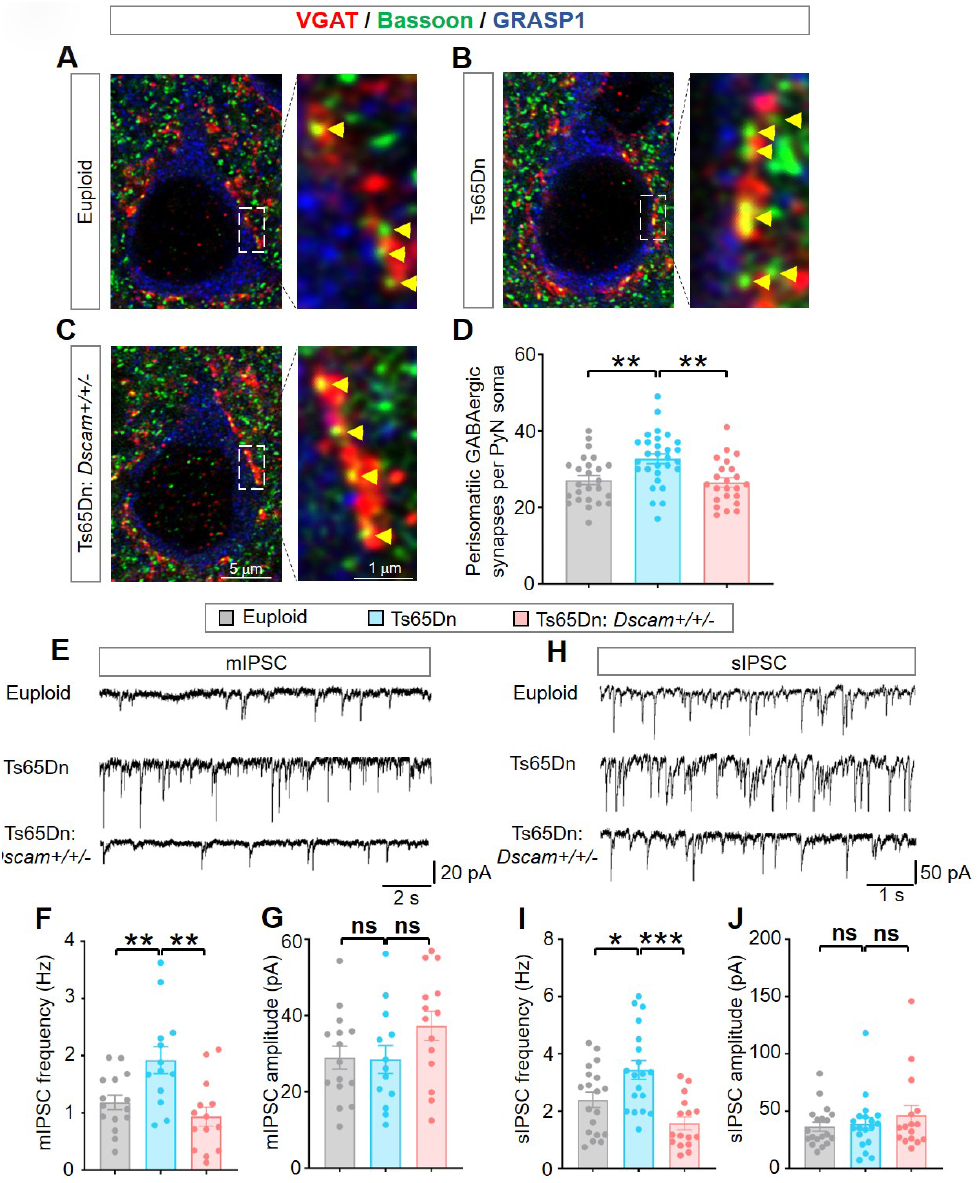
Normalizing *Dscam* levels rescues the increased synaptogenesis in basket cells and the enhanced GABAergic synaptic transmission in Ts65Dn neocortex. (A-C) Representative images of perisomatic GABAergic synapses innervating PyNs in layer II/III ACC of euploid (wild-type), Ts65Dn, and Ts65Dn:*Dscam*+/+/−. Scale bar, 5 μm. Right side is the magnified views of the regions boxed by dotted lines. Scale bar, 1 μm. The soma and proximal dendrites of PyNs were labeled by GRASP1. Yellow arrow head point to GABAergic synapses as indicated by Bassoon+ puncta that are either apposed to or overlapped with VGAT+ puncta. (D) Quantification o f t he n umber o f p erisomatic G ABAergic synapses per PyN. 5-7 PyNs were analyzed for each mouse; 4 euploid, 5 Ts65Dn, and 4 Ts65Dn:*Dscam*+/+/− mice were analyzed. Sample numbers are 24 for euploid, 29 for Ts65Dn, and 23 for Ts65Dn:*Dscam*+/+/−. (E) Representative traces of mIP-SCs from PyNs in layer II/III of ACC in euploid, Ts65Dn, and Ts65Dn:*Dscam*+/+/− brain slices. (F-G) Quantification o f m IPSC f requency (F) a nd a mplitude ( G). 24 PyNs were recorded for each mouse. 6 euploid control, 6 Ts65Dn, and 6 Ts65Dn:*Dscam*+/+/− were analyzed. N: 15 for euploid, 13 for Ts65Dn, and 14 Ts65Dn:*Dscam*+/+/−. (H) Representative traces of sIPSCs from PyNs in layer II/III of ACC in euploid, Ts65Dn and Ts65Dn:*Dscam*+/+/− brain slices. (I-J) Quantification of sIPSC frequency (I) and amplitude (J). 2 4 PyNs were recorded for each mouse. 6 euploid, 7 Ts65Dn, and 6 Ts65Dn:*Dscam*+/+/− mice were analyzed. N: 19 for euploid, 19 for Ts65Dn, and 16 Ts65Dn:*Dscam*+/+/−.

Normalizing *Dscam* levels rescues the excessive GABAergic synaptic transmission in the Ts65Dn neocortex Previous studies in Ts65Dn neocortices have demonstrated enhanced GABAergic neurogenesis (31), increased numbers of GABAergic synapses (60), and enlargement of these synapses (59). However, whether GABAergic synaptic transmission in the neocortex is increased in these mice remains to be determined. Our finding that the overexpressed *Dscam* in Ts65Dn mice increased GABAergic synaptogenesis in the neocortex prompted us to test this possibility. We examined GABAergic synaptic transmission in acute neocortical brain slices, using the whole-cell patch-clamp technique to record from PyNs in layers II/III of the ACC. We found that mIPSC frequency was increased by 63% in Ts65Dn mice compared to euploid littermates (Figures 6E-F). These results suggest an increase in presynaptic neurotransmitter release and align well with our findings of enhanced synaptogenesis in ChCs and basket cells (Figures 3E-G and 6A-D). By contrast, we found no difference in mIPSC amplitudes between euploid littermates and Ts65Dn mice (Figures 6E, G), ¬suggesting that postsynaptic responses may not be affected by the trisomy. Consistent with the role of *Dscam* in the excessive synaptogenesis in Ts65Dn mice (Figures 4G-I), normalizing *Dscam* expression prevented the increase in mIPSC frequency in these mice (Figure 6E-F), suggesting that the overexpressed *Dscam* causes excessive GABAergic synaptic transmission in the Ts65Dn neocortex. Similar changes in the frequency, but not amplitude, were observed in spontaneous inhibitory postsynaptic currents (sIPSCs) among euploid, Ts65Dn, and Ts65Dn:*Dscam*+/+/− mice (Figure 6H-J). Notably, normalizing *Dscam* levels did not rescue the increased parvalbumin-expressing (PV+) neuron density in the ACC region of Ts65Dn mice (Figures S7A-B). Consistently, the number of PV+ neurons was unaffected by loss of *Dscam* (Figures S7C-D). Thus, the overexpressed *Dscam* in the Ts65Dn neocortex causes excessive GABAergic synaptic transmission by increasing the number of GABAergic synapses without affecting the number of PV+ GABAergic neurons.

## Discussion

In this study, we provide evidence for the important role of *Dscam* expression levels in the pathogenesis of brain disorders in DS. We show that *Dscam* overexpression leads to prominent presynaptic overgrowth in ChCs and basket cells. The hyperinnervation and excessive GABAergic inhibition of PyNs in Ts65Dn were rescued by normalizing the *Dscam* levels. The converse phenotypes were observed in *Dscam* loss-of-function mutant mice. The sensitivity of GABAergic synapse development and function to *Dscam* expression levels suggest that dysregulated *Dscam* expression may underlie GABAergic dysfunction in neurological disorders that exhibit abnormal *Dscam* expression, including DS, ASDs, intractable epilepsy, bipolar disorders and, possibly, Fragile X syndrome.

### *Dscam* expression levels determine presynaptic terminal sizes in mammalian neurons

The stereotypical morphology of ChCs axon arbors and presynaptic terminals are advantageous for quantitative assessment (50, 51). Moreover, recent advances in genetic labeling of ChCs have allowed sparse labeling of single ChCs for quantifying presynaptic terminals at single-cell resolution (53, 67). By taking advantage of this system, we show in this study that *Dscam* overexpression causes presynaptic overgrowth of ChCs in the neocortex. Notably, ChC cartridge length and synaptogenesis are proportional to neocortical *Dscam* expression levels, suggesting the sensitivity of ChC development to *Dscam* levels. Thus, this work supports a conserved role of *Dscam* in regulating presynaptic terminal growth in *Drosophila* and mice. An important question is whether *Dscam* functions cell-autonomously to promote presynaptic growth in GABAergic neurons. Previous studies in *Drosophila* sensory neurons and gain-of-function studies in mouse retinal ganglion cells suggest a cell-autonomous role of *Dscam* in promoting axonal growth (11, 22). Genetic deletion of *Dscam* in single ChCs is challenging because the Cre activity in Nkx2.1-CreER mouse line is weak. Since administration of tamoxifen cannot guarantee the deletion of target genes in all or vast majority of cells in floxed mice, immunostaining or in-situ hybridization is required to confirm the deletion. However, available anti-*Dscam* antibodies are not conducive for immunostaining in neocortices (14)(data not shown). Moreover, the experimental procedures for in situ hybridization are not compatible with morphological studies of ChCs by immunostaining. Future studies with stronger Cre in ChCs will determine whether *Dscam* functions cell-autonomously.

### The coupling between presynaptic terminal growth and synaptogenesis in ChCs

Among the different types of GABAergic neurons, ChCs are thought to be the most powerful inhibitory neurons in the neocortex (45). These neurons form unique axo-axonic GABAergic synapses that selectively innervate PyNs at their AIS, where action potentials are generated (47, 48); each ChC innervates roughly two hundred PyNs (46). Impaired ChC presynaptic growth is present in individuals with epilepsy and schizophrenia, both of which are thought to be caused, in part, by disrupted GABAergic signaling (45, 68–71). Recent studies demonstrate that deleting *Erbb4*, a schizophrenia-associated gene, in ChCs causes a schizophrenia-like phenotype in mice, indicating a causal relationship between ChC defects and schizophrenia (52, 72). In addition to ErbB4, Neuregulin 1, DOCK7, Fgf13 and L1CAM have also been found to regulate the synapse formation between ChCs and PyNs (49–51, 73). In the present study, detailed investigation of the cartridge growth and synaptogenesis at single-cell resolution uncovered several interesting features of ChC development. First, presynaptic cartridge growth is strongly associated with bouton number and, to a lesser extent, bouton size (Figure 5A-D). This observation supports the synaptotropic model proposing that synaptogenesis stabilizes axonal arbor growth in neurodevelopment (74). Second, the factors that regulate cartridge growth and synaptogenesis are not necessarily the factors coupling these two processes. Although *Dscam* regulates both cartridge growth and bouton number, the coupling of these two processes remains intact in mice that are deficient of *Dscam* function, suggesting that *Dscam* is not the coupling factor. Identifying and distinguishing coupling factors from regulators is an important step toward a mechanistic understanding of ChC development.

### *Dscam* overexpression leads to excessive neocortical inhibition in DS mouse model

Overproliferation of GABAergic neurons caused by triplication of *Olig1* and *Olig2* contributes to excessive GABAergic signaling in the hippocampus (31, 32, 35). However, several lines of evidence suggest that heterogeneous etiology may exist in DS brain disorders such that different brain regions exhibit distinct molecular, cellular, and physiological defects. For example, mIPSC frequency is increased in the dentate gyrus, but not the CA1 region, of the hippocampus in adult Ts65Dn mice (31, 37). In contrast to the large body of literature on GABAergic signaling in the hippocampus of DS animal models, very little is known about whether GABAergic signaling is altered in the neocortex. Previous studies showed that the sizes of synaptic boutons and inhibitory synapses are enlarged in the neocortex of Ts65Dn mice, suggesting possible alterations in GABAergic synaptic functions in the neocortex (59, 75, 76). In the present study, we demonstrate that GABAergic inhibition of neocortical PyNs is excessive in Ts65Dn mice and that *Dscam* overexpression plays a key role in this process. Our results support the notion that alterations in GABAergic signaling in DS brains are region-specific. Normalizing *Dscam* expression reverses both increased mIPSC and sIPSC rates in neocortical PyNs (Figures 6E-J), while normalizing *Olig1* and *Olig2* expression specifically corrects s IPSC frequency in hippocampal CA1 (31). In addition, the effects of *Dscam* unlikely originate from mitotic proliferation for several reasons. Firstly, *Dscam* is expressed in differentiating neurons but not mitotic progenitors (10, 77–79). Secondly, our results show that normalizing *Dscam* levels does not rescue the increased density of parvalbumin-expressing (PV+) neurons in the ACC region of Ts65Dn mice (Figures S7A-B). Consistently, the number of cortical PV+ neurons was not affected in *Dscam* knock out mice (Figures S7C-D). Taken together, our findings suggest that the overexpressed *Dscam* in Ts65Dn mice causes presynaptic overgrowth and enhances neocortical GABAergic inhibition. They also suggest that systemic screening of hCh21 genes in *Drosophila* might be an effective approach to identify candidate genes that drive the pathogenesis of DS diseases.

### Insights into other brain disorders

Altered *Dscam* expression levels has been associated with many brain disorders, including DS, ASD, intractable epilepsy and bipolar disorder (14–19), and possibly Fragile X syndrome (11, 12, 20, 21). The present work has established a causal relationship between dysregulated *Dscam* levels and the developmental and functional defects of GABAergic neurons. The dose-dependent function of *Dscam* suggests that dysregulated *Dscam* levels may be a common pathogenic driver of GABAergic dysfunctions in related neurological diseases. For example, genetic analyses have revealed multiple disruptive single-nucleotide variants (SNVs) and copy number variants (CNVs) in *Dscam* gene in idiopathic ASD individuals, which raises the possibility of altered *Dscam* expression levels (15–17, 80). Given the regulation of GABAergic signaling by *Dscam* levels discovered in the present study and the established role of impaired GABAergic signaling in ASD (81), one may hypothesize that reduced *Dscam* expression causes impaired GABAergic signaling in ASD. In support of this, some ASD individuals with CNV of deleted enhancer region in *Dscam* show nonfebrile seizures (17), a symptom also found in mice with deficient *Dscam* function (82). It is thus important to examine *Dscam* levels in post-mortem samples of autistic individuals and determine whether altered *Dscam* expression causes GABAergic dysfunctions in ASD mouse models.

## ACKNOWLEDGEMENTS

We thank Roman Giger, Jun Wu, Andrew Nelson, Dawen Cai, Jonathan Flak, and Martin Myers for sharing reagents or technical support, Miao He, Yongjie Hou, Pedro Lowenstein, Ken Inoki, Yukiko Yamashita, and Dawen Cai for helpful discussions. This work was supported by grants from NIH (R21 NS094091 and R01 MH112669 to BY, and R37NS076752 to LLI) and Protein Folding Disease Initiative of the University of Michigan to B.Y., the Heinz C. Prechter Bipolar Research program and Richard Tam Foundation, a University of Michigan Rackham Merit Fellowship and MI-BRAIN Predoctoral Fellowship to JMH, University of Michigan Depression Center (PMJ) and the One Mind Bipolar Disorder Translational Research Award to P.M.J.

## AUTHOR CONTRIBUTIONS

H.L. and B.Y. conceived the project and designed the experiments. H.L. performed mouse breeding, drug delivery, western blotting, immunostaining, and data analysis. R.C.F, J.M.H, and G.P. performed electrophysiology recordings and analysis. T.Y., R.L. and M.V. assisted on mouse breeding and immunostaining. J.Z.H. and P.F. provided Nkx2.1-CreER and Dscam2j mice, respectively. B.Y. supervised the project. H.L., R.C.F., L.L.I., K.Y.K, P.M.J., and B.Y. wrote the paper.

---

**Fig. S1.**
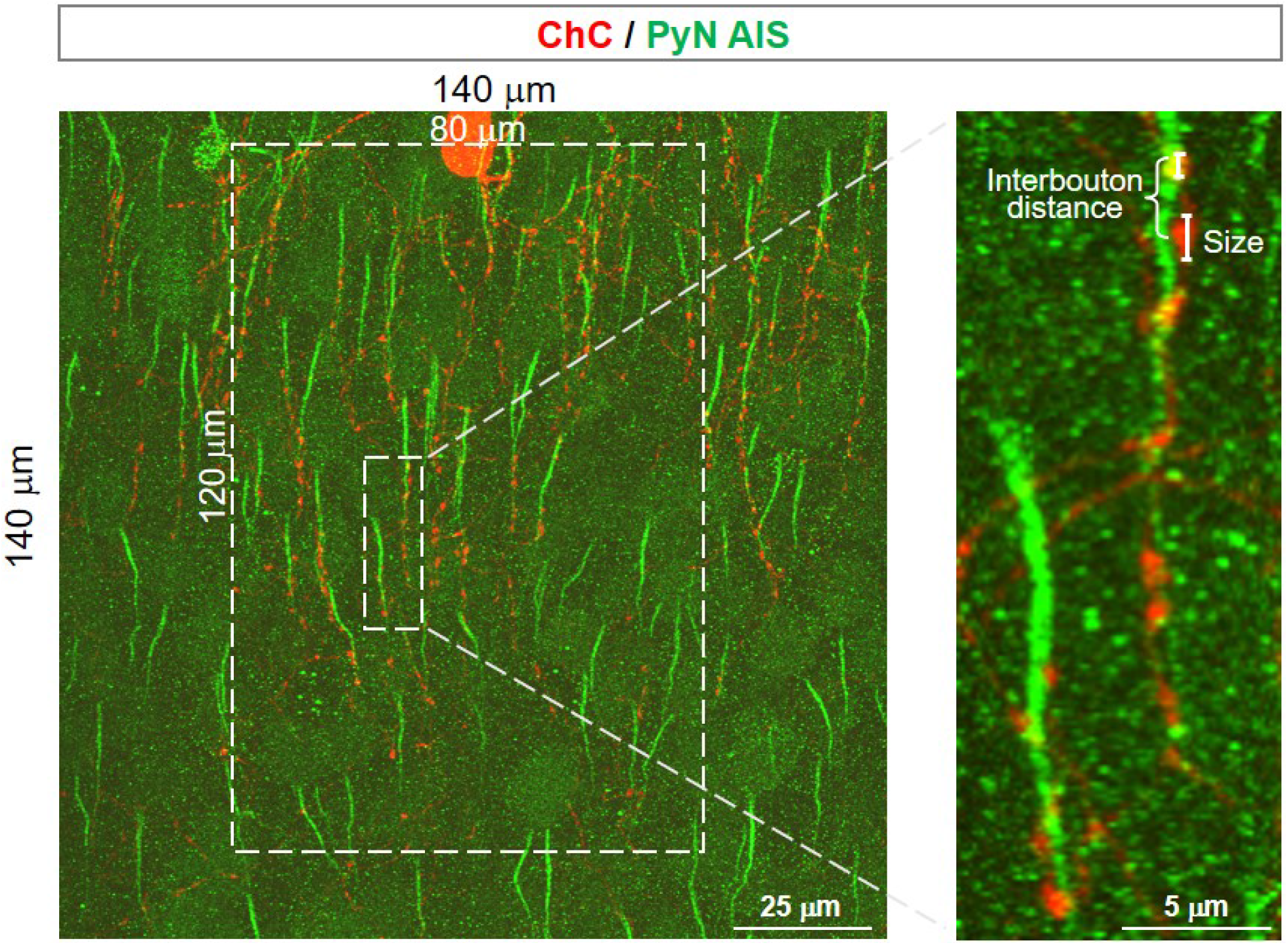
Quantification of ChC axon terminals and boutons (related to Figure 1, 3, 4). ChCs were sparsely labeled by tdTomato (red), and axon initial segments of PyNs were labeled by immunostaining with anti-phospho-IκB (Green). Confocal image stacks (0.3 μm z-steps for 100 steps) were maximally projected along the z-axis. A region of 120 μm (length) × 80 μm (width) with the cell body in the top middle was quantified. Cartridges and boutons that colocalized with AIS were quantified. Cartridge number was defined as the number of cartridges within this region. Cartridge length was defined as the distance from the first to the last bouton that colocalizes with the AIS in that cartridge. Bouton size is defined as the length of bouton in parallel to AIS. Interbouton distance is defined as the distance between two neighboring boutons.

**Fig. S2.**
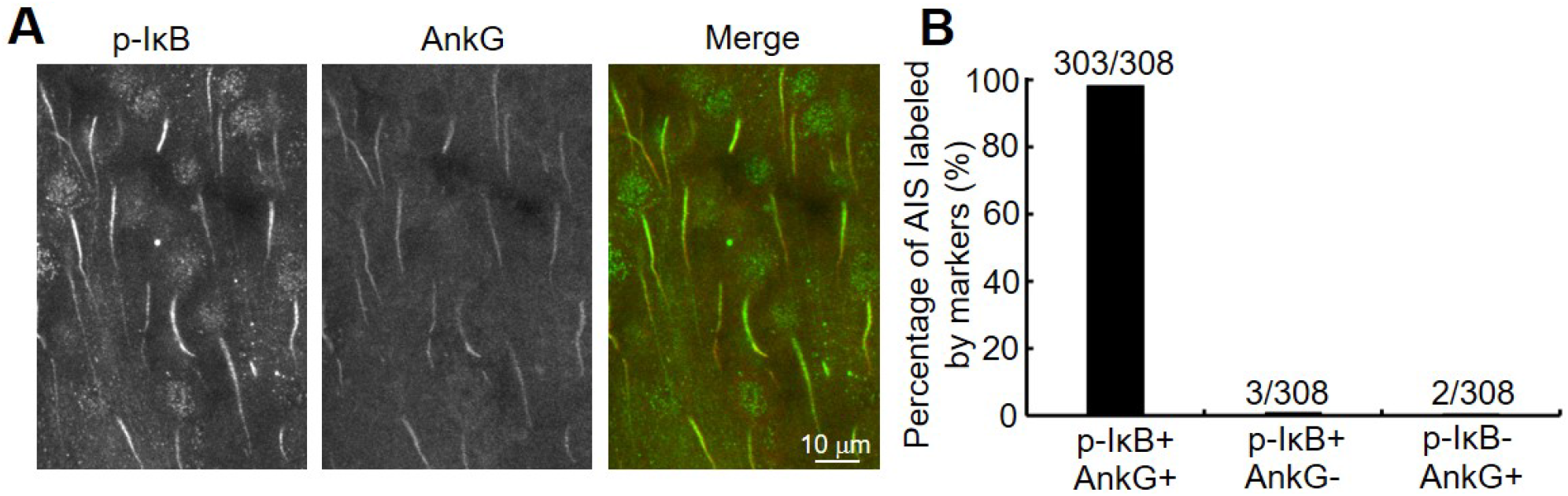
Figure 2. phospho-IκB and AnkG shows equal fidelity in labeling neocortical AIS (related to Figure 1, 3, 4). (A) AIS in layer II/III ACC was co-labeled by phospho-IκB (green) and AnkG (red). Shown are maximal projection of confocal image stacks (1 μm z-steps X 7 steps). (B) Quantification of the percentage of AIS that is labeled by phospho-IκB and/or AnkG. 308 AIS were quantified, among which 303 were co-labeled by phospho-IκB and AnkG.

**Fig. S3.**
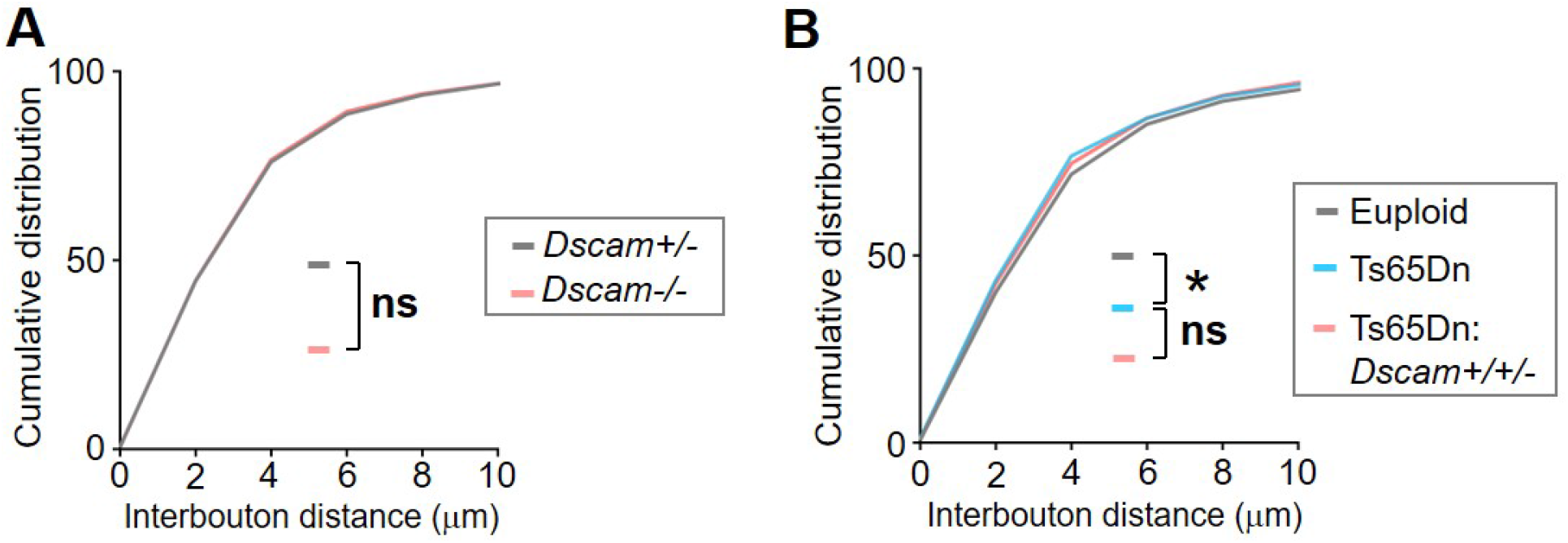
*Dscam* does not regulate bouton density (related to Figure 1, 3, 4). For each ChC, all boutons (38 194) in axon cartridges that innervate AIS in the defined volume were analyzed for bouton density (indicated by interbouton distance); 4-6 ChCs were analyzed for each mouse. The accumulative plots show the distributions of bouton density. (A) 4 *Dscam*+/− and 4 *Dscam−/−* mice were analyzed. n: 2073 for *Dscam*+/− and 1640 for *Dscam−/−*. (B) 4 euploid, 5 Ts65Dn, and 4 Ts65Dn:*Dscam*+/+/− mice were analyzed. n: 2214 for euploid, 3189 for Ts65Dn and 1912 for Ts65Dn:*Dscam*+/+/−.

**Fig. S4.**
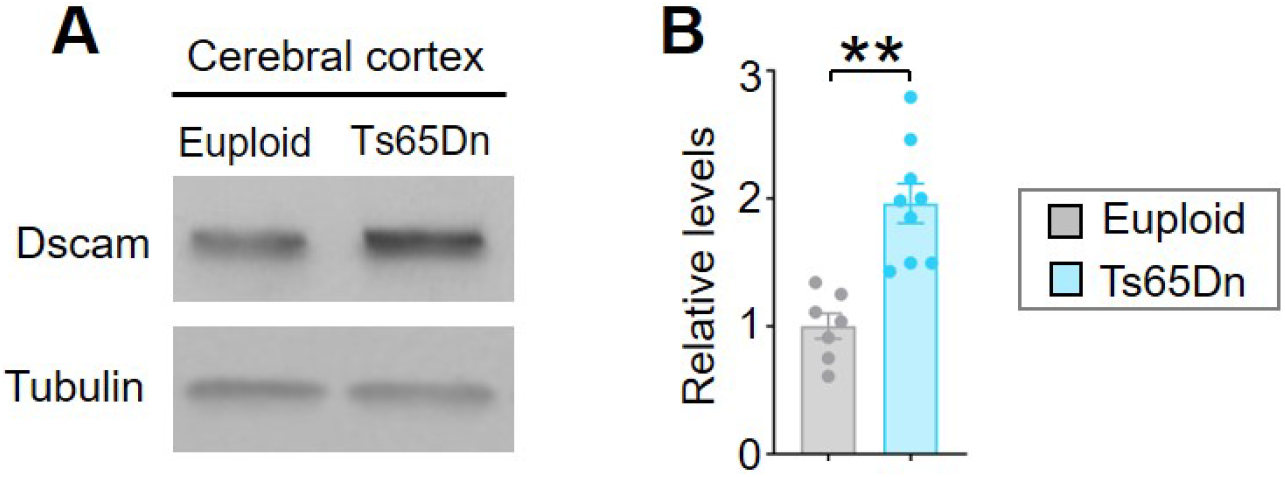
*Dscam* expression level is increased in Ts65Dn neocortices (related to Figure 3). Representative western blots (A) and quantifications (B) of neocortical samples from euploid, Ts65Dn mice. Sample numbers are n=7 for euploid, 9 for Ts65Dn.

**Fig. S5.**
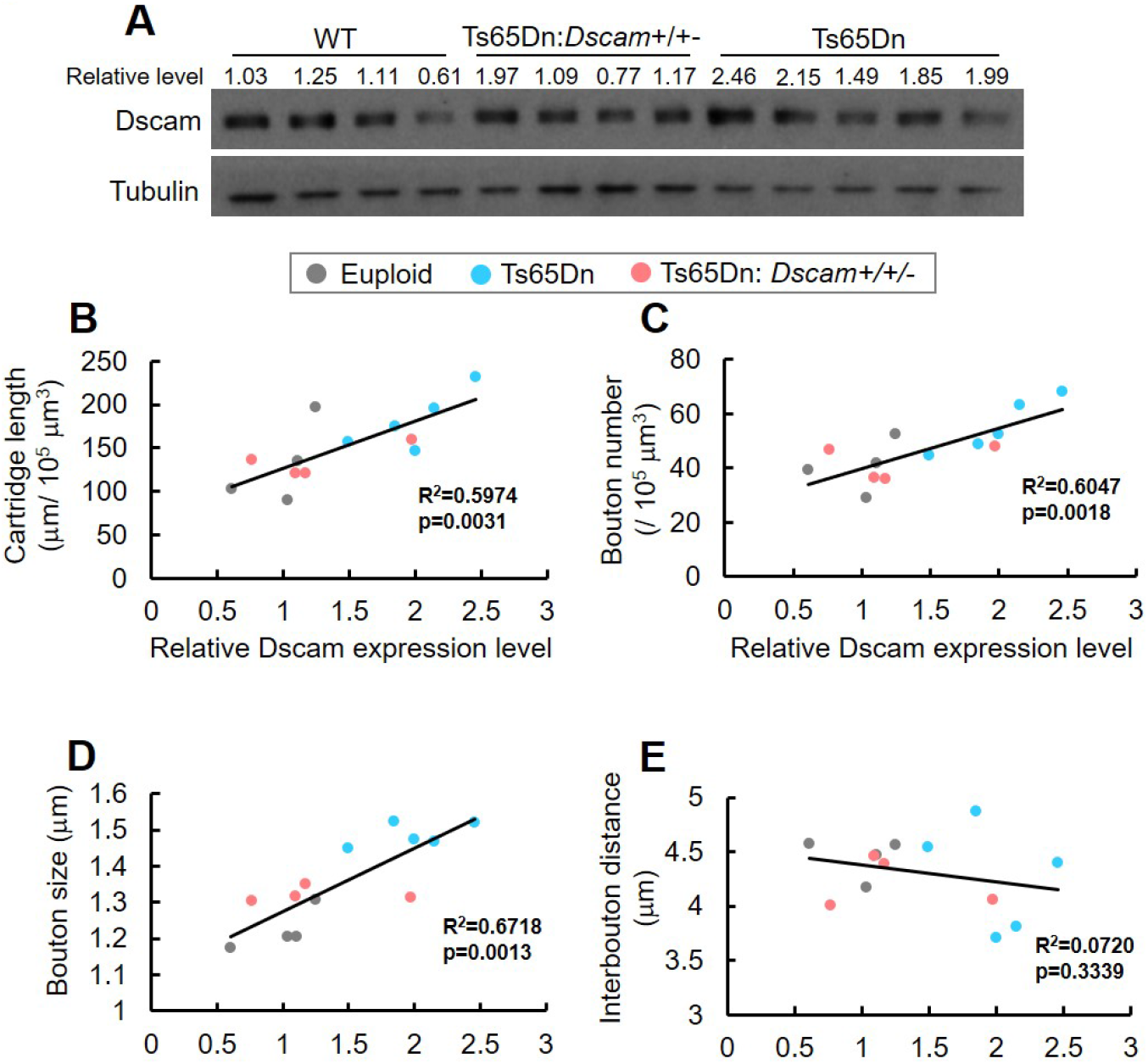
*Dscam* regulates ChC cartridge length and synaptogenesis in a dosage-dependent manner (related to Figure 5). (A) Western blots showing *Dscam* levels in the neocortex. The mouse neocortex was taken immediately after perfusion. (B-E) The correlation analyses between *Dscam* expression level and ChC cartridge length (B), bouton number (C), bouton size (D) or interbouton distance (E). In (B), the *Dscam* level of a mouse is plotted against the mean of the total cartridge length in the volume specified in Supplementary Fig. 1 in this mouse. The average cartridge length is the calculated as the average value of 4-6 ChCs sampled in each mouse. 4 euploid, 5 Ts65Dn, and 4 Ts65Dn:*Dscam*+/+/− mice were analyzed. N: 21 euploid (black dots), 26 Ts65Dn (red dots), and 21 Ts65Dn:*Dscam*+/+/− (green dots). R2 and p are calculated for linear regression.

**Fig. S6.**
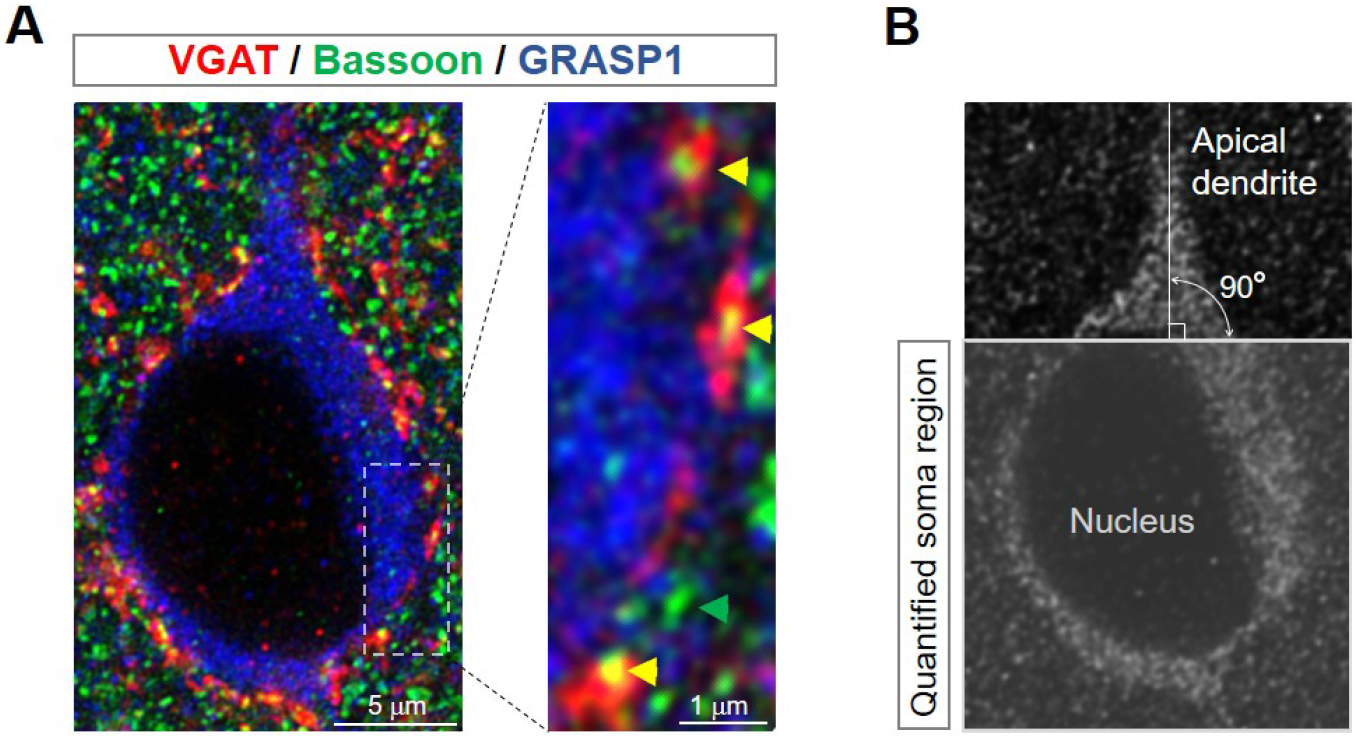
Quantification of perisomatic GABAergic synapses on PyNs (related to Figure 6). (A) Representative confocal image of a PyN in layer II/III in the ACC. The soma is labeled by anti-GRASP1 (blue). The presynaptic active zones are labeled by anti-Bassoon (green). The GABA vesicles in presynaptic terminals of GABAergic neurons are labeled by anti-VGAT (red). The perisomatic Bassoon+ puncta that are apposed to or overlap with VGAT+ puncta were quantified as GABAergic synapses (yellow arrowhead). The green arrowhead indicates the VGAT-independent perisomatic Bassoon+ puncta. (B) To define the soma region for quantifying perisomatic GABAergic synapses, we drew a line that is both perpendicular to the apical dendrite and tangent to the edge of the PyN nucleus, and then quantified GABAergic synapses in the GRASP1+ area below the line.

**Fig. S7.**
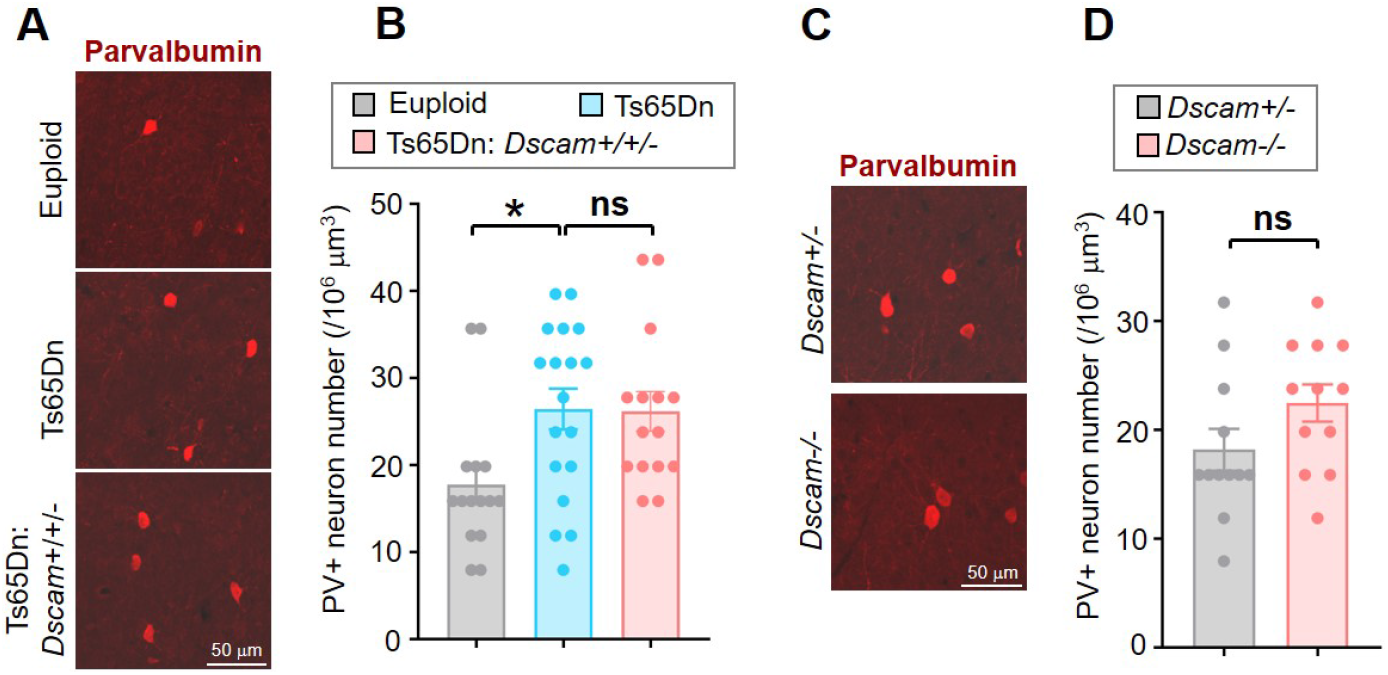
*Dscam* does not regulate the number of GABAergic neurons in the neocortex (related to Figure 6). (A-B) *Dscam* overexpression in Ts65Dn mice does not affect the number of GABAergic neurons in the neocortex. Brain sections from P28 mice were immunostained with anti-parvalbumin (PV). Representative images are shown in (A), and quantifications of the density of PV+ neurons are shown in (B). Each dot represents the value from one imaging field that is 252.4 μm (width) × 200 μm (length) × 5 μm (thickness). Images were collected from layer II/III of the ACC. Three fields in each mouse were randomly selected for imaging. 5 euploid, 6 Ts65Dn, and 5 Ts65Dn:*Dscam*+/+/− mice were analyzed. (C-D) Loss of *Dscam* does not affect the number of GABAergic neurons in the ACC. Representative images and quantifications are shown in (C) and (D), respectively. 4 *Dscam*+/− and 4 *Dscam−/−* mice were analyzed.

## Bibliography

1. L. Lim, D. Mi, A. Llorca, and O. Marin. Development and functional diversification of cortical interneurons. Neuron, 100(2):294–313, 2018. ISSN 1097-4199 (Electronic) 0896-6273 (Linking). doi: 10.1016/j.neuron.2018.10.009.

2. R. Tremblay, S. Lee, and B. Rudy. Gabaergic interneurons in the neocortex: From cellular properties to circuits. Neuron, 91(2):260–92, 2016. ISSN 1097-4199 (Electronic) 0896-6273 (xsLinking). doi: 10.1016/j.neuron.2016.06.033.

3. B. Chattopadhyaya and G. D. Cristo. Gabaergic circuit dysfunctions in neurodevelopmental disorders. Front Psychiatry, 3:51, 2012. ISSN 1664-0640 (Electronic) 1664-0640 (Linking). doi: 10.3389/fpsyt.2012.00051.

4. K. Ramamoorthi and Y. Lin. The contribution of gabaergic dysfunction to neurodevelopmental disorders. Trends Mol Med, 17(8):452–62, 2011. ISSN 1471-499X (Electronic) 1471-4914 (Linking). doi: 10.1016/j.molmed.2011.03.003.

5. M. B. Dalva, A. C. McClelland, and M. S. Kayser. Cell adhesion molecules: signalling functions at the synapse. Nature Reviews Neuroscience, 8(3):206–220, 2007. ISSN 1471-003x. doi: 10.1038/nrn2075.

6. J. de Wit and A. Ghosh. Specification of synaptic connectivity by cell surface interactions. Nat Rev Neurosci, 17(1):22–35, 2016. ISSN 1471-0048 (Electronic) 1471-003X (Linking). doi: 10.1038/nrn.2015.3.

7. J. Ko, G. Choii, and J. W. Um. The balancing act of gabaergic synapse organizers. Trends Mol Med, 21(4):256–68, 2015. ISSN 1471-499X (Electronic) 1471-4914 (Linking). doi: 10.1016/j.molmed.2015.01.004.

8. D. Krueger-Burg, T. Papadopoulos, and N. Brose. Organizers of inhibitory synapses come of age. Curr Opin Neurobiol, 45:66–77, 2017. ISSN 1873-6882 (Electronic) 0959-4388 (Linking). doi: 10.1016/j.conb.2017.04.003.

9. T. C. Sudhof. Towards an understanding of synapse formation. Neuron, 100(2):276–293, 2018. ISSN 1097-4199 (Electronic) 0896-6273 (Linking). doi: 10.1016/j.neuron.2018.09.040.

10. K. Yamakawa, Y. K. Huo, M. A. Haendel, R. Hubert, X. N. Chen, G. E. Lyons, and J. R. Korenberg. Dscam: a novel member of the immunoglobulin superfamily maps in a down syndrome region and is involved in the development of the nervous system. Human Molecular Genetics, 7(2):227–237, 1998. ISSN 0964-6906. doi: 10.1093/hmg/7.2.227.

11. J. H. Kim, X. Wang, R. Coolon, and B. Ye. Dscam expression levels determine presynaptic arbor sizes in drosophila sensory neurons. Neuron, 78(5):827–38, 2013. ISSN 1097-4199 (Electronic) 0896-6273 (Linking). doi: 10.1016/j.neuron.2013.05.020.

12. V. Cvetkovska, A. D. Hibbert, F. Emran, and B. E. Chen. Overexpression of down syndrome cell adhesion molecule impairs precise synaptic targeting. Nat Neurosci, 16(6):677–82, 2013. ISSN 1546-1726 (Electronic) 1097-6256 (Linking). doi: 10.1038/nn.3396.

13. S. A. Lowe, J. J. L. Hodge, and M. M. Usowicz. A third copy of the down syndrome cell adhesion molecule (dscam) causes synaptic and locomotor dysfunction in drosophila. Neurobiol Dis, 110:93–101, 2018. ISSN 1095-953X (Electronic) 0969-9961 (Linking). doi: 10.1016/j.nbd.2017.11.013.

14. Y. Saito, A. Oka, M. Mizuguchi, K. Motonaga, Y. Mori, L. E. Becker, K. Arima, J. Miyauchi, and S. Takashima. The developmental and aging changes of down’s syndrome cell adhesion molecule expression in normal and down’s syndrome brains. Acta neuropathologica, 100(6):654–64, 2000. ISSN 0001-6322 (Print) 0001-6322 (Linking).

15. N. Krumm, T. N. Turner, C. Baker, L. Vives, K. Mohajeri, K. Witherspoon, A. Raja, B. P. Coe, H. A. Stessman, Z. X. He, S. M. Leal, R. Bernier, and E. E. Eichler. Excess of rare, inherited truncating mutations in autism. Nat Genet, 47(6):582–8, 2015. ISSN 1546-1718 (Electronic) 1061-4036 (Linking). doi: 10.1038/ng.3303.

16. B. J. O’Roak, H. A. Stessman, E. A. Boyle, K. T. Witherspoon, B. Martin, C. Lee, L. Vives, C. Baker, J. B. Hiatt, D. A. Nickerson, R. Bernier, J. Shendure, and E. E. Eichler. Recurrent de novo mutations implicate novel genes underlying simplex autism risk. Nat Commun, 5: 5595, 2014. ISSN 2041-1723 (Electronic) 2041-1723 (Linking). doi: 10.1038/ncomms6595.

17. T. N. Turner, F. Hormozdiari, M. H. Duyzend, S. A. McClymont, P. W. Hook, I. Iossifov, A. Raja, C. Baker, K. Hoekzema, H. A. Stessman, M. C. Zody, B. J. Nelson, J. Huddleston, R. Sandstrom, J. D. Smith, D. Hanna, J. M. Swanson, E. M. Faustman, M. J. Bamshad,J. Stamatoyannopoulos, D. A. Nickerson, A. S. McCallion, R. Darnell, and E. E. Eichler. Genome sequencing of autism-affected families reveals disruption of putative noncoding regulatory dna. Am J Hum Genet, 98(1):58–74, 2016. ISSN 1537-6605 (Electronic) 0002-9297 (Linking). doi: 10.1016/j.ajhg.2015.11.023.

18. L. Shen, Z. Xiao, Y. M. Pan, M. Fang, C. S. Li, D. Chen, L. Wang, Z. Q. Xi, F. Xiao, and X. F. Wang. Altered expression of dscam in temporal lobe tissue from human and experimental animals. Synapse, 65(10):975–982, 2011. ISSN 0887-4476. doi: 10.1002/syn.20924.

19. K. Amano, K. Yamada, Y. Iwayama, S. D. Detera-Wadleigh, E. Hattori, T. Toyota, K. Tokunaga, T. Yoshikawa, and K. Yamakawa. Association study between the down syndrome cell adhesion molecule (dscam)gene and bipolar disorder. Psychiatric Genetics, 18(1):1–10, 2008. ISSN 0955-8829.

20. V. Brown, P. Jin, S. Ceman, J. C. Darnell, W. T. O’Donnell, S. A. Tenenbaum, X. Jin, Y. Feng, K. D. Wilkinson, J. D. Keene, R. B. Darnell, and S. T. Warren. Microarray identification of fmrp-associated brain mrnas and altered mrna translational profiles in fragile x syndrome. Cell, 107(4):477–87, 2001. ISSN 0092-8674 (Print) 0092-8674 (Linking).

21. J. C. Darnell, S. J. Van Driesche, C. Zhang, K. Y. Hung, A. Mele, C. E. Fraser, E. F. Stone, C. Chen, J. J. Fak, S. W. Chi, D. D. Licatalosi, J. D. Richter, and R. B. Darnell. Fmrp stalls ribosomal translocation on mrnas linked to synaptic function and autism. Cell, 146(2):247–61, 2011. ISSN 1097-4172 (Electronic) 0092-8674 (Linking). doi: 10.1016/j.cell.2011.06.013.

22. F. M. Bruce, S. Brown, J. N. Smith, P. G. Fuerst, and L. Erskine. Dscam promotes axon fasciculation and growth in the developing optic pathway. Proc Natl Acad Sci U S A, 114(7): 1702–1707, 2017. ISSN 1091-6490 (Electronic) 0027-8424 (Linking). doi: 10.1073/pnas.1618606114.

23. R. A. Santos, A. J. C. Fuertes, G. Short, K. C. Donohue, H. Shao, J. Quintanilla, P. Malakzadeh, and S. Cohen-Cory. Dscam differentially modulates pre- and postsynaptic structural and functional central connectivity during visual system wiring. Neural Dev, 13(1):22, 2018. ISSN 1749-8104 (Electronic) 1749-8104 (Linking). doi: 10.1186/s13064-018-0118-5.

24. J. Braudeau, L. Dauphinot, A. Duchon, A. Loistron, R. H. Dodd, Y. Herault, B. Delatour, and M. C. Potier. Chronic treatment with a promnesiant gaba-a alpha5-selective inverse agonist increases immediate early genes expression during memory processing in mice and rectifies their expression levels in a down syndrome mouse model. Adv Pharmacol Sci, 2011:153218, 2011. ISSN 1687-6342 (Electronic) 1687-6334 (Linking). doi: 10.1155/2011/153218.

25. J. Braudeau, B. Delatour, A. Duchon, P. L. Pereira, L. Dauphinot, F. de Chaumont, J. C. Olivo-Marin, R. H. Dodd, Y. Herault, and M. C. Potier. Specific targeting of the gaba-a receptor alpha5 subtype by a selective inverse agonist restores cognitive deficits in down syndrome mice. J Psychopharmacol, 25(8):1030–42, 2011. ISSN 1461-7285 (Electronic) 0269-8811 (Linking). doi: 10.1177/0269881111405366.

26. D. Colas, B. Chuluun, D. Warrier, M. Blank, D. Z. Wetmore, P. Buckmaster, C. C. Garner, and H. C. Heller. Short-term treatment with the gabaa receptor antagonist pentylenetetrazole produces a sustained pro-cognitive benefit in a mouse model of down’s syndrome. British Journal of Pharmacology, 169(5):963–973, 2013. ISSN 0007-1188. doi: 10.1111/bph.12169.

27. F. Fernandez, W. Morishita, E. Zuniga, J. Nguyen, M. Blank, R. C. Malenka, and C. C. Garner. Pharmacotherapy for cognitive impairment in a mouse model of down syndrome. Nat Neurosci, 10(4):411–3, 2007. ISSN 1097-6256 (Print) 1097-6256 (Linking). doi: 10.1038/nn1860.

28. C. Martinez-Cue, P. Martinez, N. Rueda, R. Vidal, S. Garcia, V. Vidal, A. Corrales, J. A. Montero, A. Pazos, J. Florez, R. Gasser, A. W. Thomas, M. Honer, F. Knoflach, J. L. Trejo, J. G. Wettstein, and M. C. Hernandez. Reducing gabaa alpha5 receptor-mediated inhibition rescues functional and neuromorphological deficits in a mouse model of down syndrome. J Neurosci, 33(9):3953–66, 2013. ISSN 1529-2401 (Electronic) 0270-6474 (Linking). doi: 10.1523/JNEUROSCI.1203-12.2013.

29. R. H. Reeves, N. G. Irving, T. H. Moran, A. Wohn, C. Kitt, S. S. Sisodia, C. Schmidt, R. T. Bronson, and M. T. Davisson. A mouse model for down syndrome exhibits learning and behaviour deficits. Nat Genet, 11(2):177–84, 1995. ISSN 1061-4036 (Print) 1061-4036 (Linking). doi: 10.1038/ng1095-177.

30. N. Rueda, J. Florez, and C. Martinez-Cue. Mouse models of down syndrome as a tool to unravel the causes of mental disabilities. Neural Plasticity, 2012. ISSN 2090-5904. doi: Artn58407110.1155/2012/584071.

31. L. Chakrabarti, T. K. Best, N. P. Cramer, R. S. Carney, J. T. Isaac, Z. Galdzicki, and T. F. Haydar. Olig1 and olig2 triplication causes developmental brain defects in down syndrome. Nat Neurosci, 13(8):927–34, 2010. ISSN 1546-1726 (Electronic) 1097-6256 (Linking). doi: 10.1038/nn.2600.

32. A. Contestabile, S. Magara, and L. Cancedda. The gabaergic hypothesis for cognitive disabilities in down syndrome. Front Cell Neurosci, 11:54, 2017. ISSN 1662-5102 (Print) 1662-5102 (Linking). doi: 10.3389/fncel.2017.00054.

33. A. C. Costa and M. J. Grybko. Deficits in hippocampal ca1 ltp induced by tbs but not hfs in the ts65dn mouse: a model of down syndrome. Neurosci Lett, 382(3):317–22, 2005. ISSN 0304-3940 (Print) 0304-3940 (Linking). doi: 10.1016/j.neulet.2005.03.031.

34. F. Fernandez and C. C. Garner. Episodic-like memory in ts65dn, a mouse model of down syndrome. Behav Brain Res, 188(1):233–7, 2008. ISSN 0166-4328 (Print) 0166-4328 (Linking). doi: 10.1016/j.bbr.2007.09.015.

35. T. F. Haydar and R. H. Reeves. Trisomy 21 and early brain development. Trends Neurosci, 35(2):81–91, 2012. ISSN 1878-108X (Electronic) 0166-2236 (Linking). doi: 10.1016/j.tins.2011.11.001.

36. A. M. Kleschevnikov, P. V. Belichenko, M. Faizi, L. F. Jacobs, K. Htun, M. Shamloo, and W. C. Mobley. Deficits in cognition and synaptic plasticity in a mouse model of down syndrome ameliorated by gabab receptor antagonists. J Neurosci, 32(27):9217–27, 2012. ISSN 1529-2401 (Electronic) 0270-6474 (Linking). doi: 10.1523/JNEUROSCI.1673-12.2012.

37. A. M. Kleschevnikov, P. V. Belichenko, J. Gall, L. George, R. Nosheny, M. T. Maloney, A. Salehi, and W. C. Mobley. Increased efficiency of the gabaa and gabab receptor-mediated neurotransmission in the ts65dn mouse model of down syndrome. Neurobiol Dis, 45(2):683–91, 2012. ISSN 1095-953X (Electronic) 0969-9961 (Linking). doi: 10.1016/j.nbd.2011.10.009.

38. A. M. Kleschevnikov, P. V. Belichenko, A. J. Villar, C. J. Epstein, R. C. Malenka, and W. C. Mobley. Hippocampal long-term potentiation suppressed by increased inhibition in the ts65dn mouse, a genetic model of down syndrome. J Neurosci, 24(37):8153–60, 2004. ISSN 1529-2401 (Electronic) 0270-6474 (Linking). doi: 10.1523/JNEUROSCI.1766-04.2004.

39. R. J. Siarey, J. Stoll, S. I. Rapoport, and Z. Galdzicki. Altered long-term potentiation in the young and old ts65dn mouse, a model for down syndrome. Neuropharmacology, 36(11-12): 1549–54, 1997. ISSN 0028-3908 (Print) 0028-3908 (Linking).

40. B. Yang, J. B. Treweek, R. P. Kulkarni, B. E. Deverman, C. K. Chen, E. Lubeck, S. Shah, L. Cai, and V. Gradinaru. Single-cell phenotyping within transparent intact tissue through whole-body clearing. Cell, 158(4):945–958, 2014. ISSN 0092-8674. doi: 10.1016/j.cell.2014.07.017.

41. A. D. Nelson, R. N. Caballero-Floran, J. C. R. Diaz, J. M. Hull, Y. Yuan, J. Li, K. Chen, K. K. Walder, L. F. Lopez-Santiago, V. Bennett, M. G. McInnis, L. L. Isom, C. Wang, M. Zhang, K. S. Jones, and P. M. Jenkins. Correction: Ankyrin-g regulates forebrain connectivity and network synchronization via interaction with gabarap. Mol Psychiatry, 2019. ISSN 1476-5578 (Electronic) 1359-4184 (Linking). doi: 10.1038/s41380-019-0361-0.

42. B. Ye, D. Liao, X. Zhang, P. Zhang, H. Dong, and R. L. Huganir. Grasp-1: a neuronal rasgef associated with the ampa receptor/grip complex. Neuron, 26(3):603–17, 2000.

43. J. N. Flak, D. Arble, W. Pan, C. Patterson, T. Lanigan, P. B. Goforth, J. Sacksner, M. Joosten, D. A. Morgan, M. B. Allison, J. Hayes, E. Feldman, R. J. Seeley, D. P. Olson, K. Rahmouni, and M. G. Myers. A leptin-regulated circuit controls glucose mobilization during noxious stimuli. Journal of Clinical Investigation, 127(8):3103–3113, 2017. ISSN 0021-9738. doi: 10.1172/Jci90147.

44. A. D. Nelson, R. N. Caballero-Floran, J. C. Rodriguez Diaz, J. M. Hull, Y. Yuan, J. Li, K. Chen, K. K. Walder, L. F. Lopez-Santiago, V. Bennett, M. G. McInnis, L. L. Isom, C. Wang, M. Zhang, K. S. Jones, and P. M. Jenkins. Ankyrin-g regulates forebrain connectivity and network synchronization via interaction with gabarap. Mol Psychiatry, 2018. ISSN 1476-5578 (Electronic) 1359-4184 (Linking). doi: 10.1038/s41380-018-0308-x.

45. J. DeFelipe. Chandelier cells and epilepsy. Brain, 122 (Pt 10):1807–22, 1999. ISSN 0006-8950 (Print) 0006-8950 (Linking).

46. L. Blazquez-Llorca, A. Woodruff, M. Inan, S. A. Anderson, R. Yuste, J. DeFelipe, and A. Merchan-Perez. Spatial distribution of neurons innervated by chandelier cells. Brain Struct Funct, 220(5):2817–34, 2015. ISSN 1863-2661 (Electronic) 1863-2653 (Linking). doi: 10.1007/s00429-014-0828-3.

47. E. G. Jones. Varieties and distribution of non-pyramidal cells in the somatic sensory cortex of the squirrel monkey. J Comp Neurol, 160(2):205–67, 1975. ISSN 0021-9967 (Print) 0021-9967 (Linking). doi: 10.1002/cne.901600204.

48. J. Szentagothai. Module-concept in cerebral-cortex architecture. Brain Research, 95(2-3): 475–496, 1975. ISSN 0006-8993. doi: Doi10.1016/0006-8993(75)90122-5.

49. E. Favuzzi, R. Deogracias, A. Marques-Smith, P. Maeso, J. Jezequel, D. Exposito-Alonso, M. Balia, T. Kroon, A. J. Hinojosa, F. Maraver E, and B. Rico. Distinct molecular programs regulate synapse specificity in cortical inhibitory circuits. Science, 363(6425):413–417, 2019. ISSN 1095-9203 (Electronic) 0036-8075 (Linking). doi: 10.1126/science.aau8977.

50. P. Fazzari, A. V. Paternain, M. Valiente, R. Pla, R. Lujan, K. Lloyd, J. Lerma, O. Marin, and B. Rico. Control of cortical gaba circuitry development by nrg1 and erbb4 signalling. Nature, 464(7293):1376–80, 2010. ISSN 1476-4687 (Electronic) 0028-0836 (Linking). doi: 10.1038/nature08928.

51. Y. Tai, J. A. Janas, C. L. Wang, and L. Van Aelst. Regulation of chandelier cell cartridge and bouton development via dock7-mediated erbb4 activation. Cell Rep, 6(2):254–63, 2014. ISSN 2211-1247 (Electronic). doi: 10.1016/j.celrep.2013.12.034.

52. J. M. Yang, C. J. Shen, X. J. Chen, Y. Kong, Y. S. Liu, X. W. Li, Z. Chen, T. M. Gao, and X. M. Li. erbb4 deficits in chandelier cells of the medial prefrontal cortex confer cognitive dysfunctions: Implications for schizophrenia. Cereb Cortex, 2018. ISSN 1460-2199 (Electronic) 1047-3211 (Linking). doi: 10.1093/cercor/bhy316.

53. H. Taniguchi, J. Lu, and Z. J. Huang. The spatial and temporal origin of chandelier cells in mouse neocortex. Science, 339(6115):70–4, 2013. ISSN 1095-9203 (Electronic) 0036-8075 (Linking). doi: 10.1126/science.1227622.

54. M. Inan and S. A. Anderson. The chandelier cell, form and function. Curr Opin Neurobiol, 26:142–8, 2014. ISSN 1873-6882 (Electronic) 0959-4388 (Linking). doi: 10.1016/j.conb.2014.01.009.

55. A. Steinecke, E. Hozhabri, S. Tapanes, Y. Ishino, H. Zeng, N. Kamasawa, and H. Taniguchi. Neocortical chandelier cells developmentally shape axonal arbors through reorganization but establish subcellular synapse specificity without refinement. Eneuro, 4(3), 2017. ISSN 2373-2822. doi: UNSPe005710.1523/ENEURO.0057-17.2017.

56. J. Lu, J. Tucciarone, N. Padilla-Coreano, M. He, J. A. Gordon, and Z. J. Huang. Selective inhibitory control of pyramidal neuron ensembles and cortical subnetworks by chandelier cells. Nat Neurosci, 20(10):1377–1383, 2017. ISSN 1546-1726 (Electronic) 1097-6256 (Linking). doi: 10.1038/nn.4624.

57. S. A. Buffington, J. M. Sobotzik, C. Schultz, and M. N. Rasband. I kappa b alpha is not required for axon initial segment assembly. Molecular and Cellular Neuroscience, 50(1): 1–9, 2012. ISSN 1044-7431. doi: 10.1016/j.mcn.2012.03.003.

58. D. G. Herrera and H. A. Robertson. Activation of c-fos in the brain. Prog Neurobiol, 50(2-3): 83–107, 1996. ISSN 0301-0082 (Print) 0301-0082 (Linking).

59. M. A. Kurt, D. C. Davies, M. Kidd, M. Dierssen, and J. Florez. Synaptic deficit in the temporal cortex of partial trisomy 16 (ts65dn) mice. Brain Res, 858(1):191–7, 2000. ISSN 0006-8993 (Print) 0006-8993 (Linking).

60. R. L. Nosheny, P. V. Belichenko, B. L. Busse, A. M. Weissmiller, V. Dang, D. Das, A. Fahimi, A. Salehi, S. J. Smith, and W. C. Mobley. Increased cortical synaptic activation of trkb and downstream signaling markers in a mouse model of down syndrome. Neurobiol Dis, 77: 173–90, 2015. ISSN 1095-953X (Electronic) 0969-9961 (Linking). doi: 10.1016/j.nbd.2015.02.022.

61. Gabriella R Sterne, Jung Hwan Kim, and Bing Ye. Dysregulated dscam levels act through abelson tyrosine kinase to enlarge presynaptic arbors. eLife, 4:e05196, 2015.

62. A. Javaherian and H. T. Cline. Coordinated motor neuron axon growth and neuromuscular synaptogenesis are promoted by cpg15 in vivo. Neuron, 45(4):505–12, 2005. ISSN 0896-6273 (Print) 0896-6273 (Linking). doi: 10.1016/j.neuron.2004.12.051.

63. A. Paul, M. Crow, R. Raudales, M. He, J. Gillis, and Z. J. Huang. Transcriptional architecture of synaptic communication delineates gabaergic neuron identity. Cell, 171(3):522–539 e20, 2017. ISSN 1097-4172 (Electronic) 0092-8674 (Linking). doi: 10.1016/j.cell.2017.08.032.

64. Y. Wang, A. Gupta, M. Toledo-Rodriguez, C. Z. Wu, and H. Markram. Anatomical, physiological, molecular and circuit properties of nest basket cells in the developing somatosensory cortex. Cereb Cortex, 12(4):395–410, 2002. ISSN 1047-3211 (Print) 1047-3211 (Linking). doi: 10.1093/cercor/12.4.395.

65. S. T. Dieck, L. Sanmarti-Vila, K. Langnaese, K. Richter, S. Kindler, A. Soyke, H. Wex, K. H. Smalla, U. Kampf, J. T. Franzer, M. Stumm, C. C. Garner, and E. D. Gundelfinger. Bassoon, a novel zinc-finger cag/glutamine-repeat protein selectively localized at the active zone of presynaptic nerve terminals. Journal of Cell Biology, 142(2):499–509, 1998. ISSN 0021-9525. doi: DOI10.1083/jcb.142.2.499.

66. F. A. Chaudhry, R. J. Reimer, E. E. Bellocchio, N. C. Danbolt, K. K. Osen, R. H. Edwards, and J. Storm-Mathisen. The vesicular gaba transporter, vgat, localizes to synaptic vesicles in sets of glycinergic as well as gabaergic neurons. Journal of Neuroscience, 18(23):9733– 9750, 1998. ISSN 0270-6474.

67. X. Wang, J. Tucciarone, S. Jiang, F. Yin, B. S. Wang, D. Wang, Y. Jia, X. Jia, Y. Li, T. Yang, Z. Xu, M. A. Akram, Y. Wang, S. Zeng, G. A. Ascoli, P. Mitra, H. Gong, Q. Luo, and Z. J. Huang. Genetic single neuron anatomy reveals fine granularity of cortical axo-axonic cells. Cell Rep, 26(11):3145–3159 e5, 2019. ISSN 2211-1247 (Electronic). doi: 10.1016/j.celrep.2019.02.040.

68. D. A. Lewis. Inhibitory neurons in human cortical circuits: substrate for cognitive dysfunction in schizophrenia. Curr Opin Neurobiol, 26:22–6, 2014. ISSN 1873-6882 (Electronic) 0959-4388 (Linking). doi: 10.1016/j.conb.2013.11.003.

69. D. A. Lewis, A. A. Curley, J. R. Glausier, and D. W. Volk. Cortical parvalbumin interneurons and cognitive dysfunction in schizophrenia. Trends Neurosci, 35(1):57–67, 2012. ISSN 1878-108X (Electronic) 0166-2236 (Linking). doi: 10.1016/j.tins.2011.10.004.

70. K. Nakazawa, V. Zsiros, Z. Jiang, K. Nakao, S. Kolata, S. Zhang, and J. E. Belforte. Gabaergic interneuron origin of schizophrenia pathophysiology. Neuropharmacology, 62(3):1574– 83, 2012. ISSN 1873-7064 (Electronic) 0028-3908 (Linking). doi: 10.1016/j.neuropharm.2011.01.022.

71. T. U. Woo, R. E. Whitehead, D. S. Melchitzky, and D. A. Lewis. A subclass of prefrontal gamma-aminobutyric acid axon terminals are selectively altered in schizophrenia. Proc Natl Acad Sci U S A, 95(9):5341–6, 1998. ISSN 0027-8424 (Print) 0027-8424 (Linking).

72. I. Del Pino, C. Garcia-Frigola, N. Dehorter, J. R. Brotons-Mas, E. Alvarez-Salvado, M. Martinez de Lagran, G. Ciceri, M. V. Gabaldon, D. Moratal, M. Dierssen, S. Canals, O. Marin, and B. Rico. Erbb4 deletion from fast-spiking interneurons causes schizophrenia-like phenotypes. Neuron, 79(6):1152–68, 2013. ISSN 1097-4199 (Electronic) 0896-6273 (Linking). doi: 10.1016/j.neuron.2013.07.010.

73. Y. Tai, N. B. Gallo, M. Wang, J. R. Yu, and L. Van Aelst. Axo-axonic innervation of neocortical pyramidal neurons by gabaergic chandelier cells requires ankyring-associated l1cam. Neuron, 102(2):358–372 e9, 2019. ISSN 1097-4199 (Electronic) 0896-6273 (Linking). doi: 10.1016/j.neuron.2019.02.009.

74. Hollis Cline and Kurt Haas. The regulation of dendritic arbor development and plasticity by glutamatergic synaptic input: a review of the synaptotrophic hypothesis. J Physiol, 586(6): 1509–1517, 2008. doi: 10.1113/jphysiol.2007.150029.

75. P. V. Belichenko, A. M. Kleschevnikov, E. Masliah, C. Wu, R. Takimoto-Kimura, A. Salehi, and W. C. Mobley. Excitatory-inhibitory relationship in the fascia dentata in the ts65dn mouse model of down syndrome. J Comp Neurol, 512(4):453–66, 2009. ISSN 1096-9861 (Electronic) 0021-9967 (Linking). doi: 10.1002/cne.21895.

76. P. V. Belichenko, E. Masliah, A. M. Kleschevnikov, A. J. Villar, C. J. Epstein, A. Salehi, and W. C. Mobley. Synaptic structural abnormalities in the ts65dn mouse model of down syndrome. J Comp Neurol, 480(3):281–98, 2004. ISSN 0021-9967 (Print) 0021-9967 (Linking). doi: 10.1002/cne.20337.

77. D. D. Larsen and E. M. Callaway. Development of layer-specific axonal arborizations in mouse primary somatosensory cortex. J Comp Neurol, 494(3):398–414, 2006. ISSN 0021-9967 (Print) 0021-9967 (Linking). doi: 10.1002/cne.20754.

78. C. Mayer, C. Hafemeister, R. C. Bandler, R. Machold, R. Batista Brito, X. Jaglin, K. Allaway, A. Butler, G. Fishell, and R. Satija. Developmental diversification of cortical inhibitory interneurons. Nature, 555(7697):457–462, 2018. ISSN 1476-4687 (Electronic) 0028-0836 (Linking). doi: 10.1038/nature25999.

79. D. Mi, Z. Li, L. Lim, M. Li, M. Moissidis, Y. Yang, T. Gao, T. X. Hu, T. Pratt, D. J. Price, N. Sestan, and O. Marin. Early emergence of cortical interneuron diversity in the mouse embryo. Science, 360(6384):81–85, 2018. ISSN 1095-9203 (Electronic) 0036-8075 (Linking). doi: 10.1126/science.aar6821.

80. T. Y. Wang, H. Guo, B. Xiong, H. A. F. Stessman, H. D. Wu, B. P. Coe, T. N. Turner, Y. L. Liu, W. J. Zhao, K. Hoekzema, L. Vives, L. Xia, M. N. Tang, J. J. Ou, B. Y. Chen, Y. D. Shen, G. L. Xun, M. Long, J. Lin, Z. N. Kronenberg, Y. Peng, T. Bai, H. H. Li, X. Y. Ke, Z. M. Hu, J. P. Zhao, X. B. Zou, K. Xia, and E. E. Eichler. De novo genic mutations among a chinese autism spectrum disorder cohort. Nature Communications, 7, 2016. ISSN 2041-1723. doi: ARTN1331610.1038/ncomms13316.

81. G. Cellot and E. Cherubini. Gabaergic signaling as therapeutic target for autism spectrum disorders. Frontiers in Pediatrics, 2, 2014. ISSN 2296-2360. doi: ARTN7010.3389/fped. 2014.00070.

82. P. G. Fuerst, A. Koizumi, R. H. Masland, and R. W. Burgess. Neurite arborization and mosaic spacing in the mouse retina require dscam. Nature, 451(7177):470–U8, 2008. ISSN 0028-0836. doi: 10.1038/nature06514.

